# ACSS2 regulates ferroptosis in an E2F1-dependent manner in breast cancer brain metastatic cells

**DOI:** 10.1101/2024.10.18.619082

**Authors:** Emily M. Esquea, Riley G. Young, Lorela Ciraku, Jessica Merzy, Nusaiba N. Ahmed, Alexandra N. Talarico, Mangalam Karuppiah, Wiktoria Gocal, Nicole L. Simone, Alexej Dick, Mauricio J. Reginato

**Affiliations:** Department of Biochemistry and Molecular Biology, Drexel University College of Medicine, Philadelphia, PA 19102; Department of Radiation Oncology, Sidney Kimmel Cancer Center, Thomas Jefferson University, Philadelphia, PA; Cancer Risk and Control Program, Thomas Jefferson University, Philadelphia, PA 19107, USA; Translational and Cellular Oncology Program, Sidney Kimmel Cancer Center, Thomas Jefferson University, Philadelphia, PA 19107, USA

**Keywords:** O-GlcNAc, OGT, hexosamine, cancer, metabolism, breast cancer, brain metastasis, acetate, acetyl-CoA, ACSS2, CDK5, lipids, ferroptosis, E2F1, SLC7A11, GPX4

## Abstract

Brain metastasis diagnosis in breast cancer patients is considered an end-stage event. The median survival after diagnosis is measured in months, thus there is an urgent need to develop novel treatment strategies. Breast cancers that metastasize to the brain must adapt to the unique brain environment and are highly dependent on acetate metabolism for growth and survival. However, the signaling pathways that regulate survival in breast cancer brain metastatic (BCBM) tumors are not known. Primary brain tumor cells can convert acetate to acetyl-CoA via phosphorylation of acetyl-CoA synthetase 2 (ACSS2) by the cyclin-dependent kinase-5 (CDK5) regulated by the nutrient sensor O-GlcNAc transferase (OGT). Here, we show that breast cancer cells selected to metastasize to the brain contain increased levels of O-GlcNAc, OGT and ACSS2-Ser267 phosphorylation compared to parental breast cancer cells. Moreover, OGT and CDK5 are required for breast cancer cell growth in the brain parenchyma *in vivo.* Importantly, ACSS2 and ACSS2-S267D phospho-mimetic mutant are critical for *in vivo* breast cancer growth in the brain but not in the mammary fat pad. Mechanistically, we show that ACSS2 regulates BCBM cell survival by suppressing ferroptosis via regulation of E2F1-mediated expression of anti-ferroptotic proteins SLC7A11 and GPX4. Lastly, we show treatment with a novel brain-permeable small molecule ACSS2 inhibitor induced ferroptosis and reduced BCBM growth *ex vivo* and *in vivo*. These results suggest a crucial role for ACSS2 in protecting from ferroptosis in breast cancer brain metastatic cells and suggests that breast cancer brain metastatic cells may be susceptible to ferroptotic inducers.

## INTRODUCTION

Breast cancer brain metastases (BCBM) remains a devastating consequence of breast cancer progression, contributing to reduced overall survival post-diagnosis. Breast cancer is the second most common cancer to metastasize to the brain, with the triple-negative breast cancer (TNBC) subtype exhibiting the highest incidence rate compared to other molecular subtypes [1]. TNBC brain metastases has the lowest overall survival (OS) ranging from two months in untreated cases to 4.9 months [1] [2], highlighting an urgent need to understand the molecular mechanisms of growth and survival in the brain microenvironment to aid in developing targeted therapies.

The brain provides a unique metabolic microenvironment. Glucose is the primary oxidative fuel for the healthy brain [3] [4] [5], while other metabolites such as glutamine and acetate are recognized as preferred fuels for astrocytes [6] [4]. The blood-brain barrier (BBB) limits nutrient availability to the brain creating an environment that is scarce of metabolites, growth factors, proteins, and amino acids [7] [8] [9]. Importantly, the cerebrospinal fluid and brain tissue contain low levels of lipids [10] and triacylglycerides [11] compared to other tissues, reflecting a distinct physiological feature that imposes a specific requirement for metabolic adaptation to this microenvironment. *In situ* ^13^C-glucose infusion studies revealed that although tumors in patients with glioblastoma (GBM), lung, and breast cancer brain metastases oxidize glucose in the citric acid cycle, a significant portion of the acetyl-CoA pool was derived from other metabolic sources, including acetate [12] [13]. Outside the gut, the brain has the highest levels of acetate [14], and acetate has been shown to support the growth of tumors in the brain [13]. These observations were accompanied by evidence of upregulation of the enzyme that catalyzes the conversion of acetate to acetyl-CoA, acetyl-CoA synthetase 2 (ACSS2), in these tumors, which has been shown to play a key role in GBM [13] [15] [16]. Acetate uptake and upregulation of ACSS2 creates an increase in the acetyl-CoA pool, which can serve as a substrate for *de novo* lipid biosynthesis enzymes and histone acetylation, ultimately regulating gene expression that support growth and survival in cancer cells [17]. However, whether ACSS2 regulates growth in breast cancer brain metastatic cells is not known.

UDP-N-acetylglucosamine (UDP-GlcNAc) is generated by the hexosamine biosynthetic pathway, which utilizes major metabolites [18]. UDP-GlcNAc serves as a substrate for N- and O-Glycosylation, regulating cellular behaviors in response to nutrient availability [19]. It also acts as a substrate for O-linked ß-N-acetylglucosamine (O-GlcNAc) transferase (OGT), which catalyzes the addition of O-GlcNAc moieties onto serine and threonine residues of nuclear and cytoplasmic proteins [20]. O-GlcNAcylation is a post-translational modification that alters protein functions, protein-protein interactions, and phosphorylation states [21]. OGT and O-GlcNAcylation are elevated in most cancers [22], including breast cancer and glioblastoma (GBM), and targeting this modification inhibits tumor cell growth *in vitro* and *in vivo* [23], [24]. We have previously shown that OGT is upregulated and required for glioblastoma growth [25]. Elevated expression of OGT promotes phosphorylation of ACSS2 at Ser267 (p-Ser267-ACSS2) via a non-canonical cyclin-dependent kinase 5 (CDK5). This post-translational modification leads to increased stability of the ACSS2 protein and promotes increased acetyl-CoA levels [16]. Given the metabolic similarities between GBM and brain metastases [16], we hypothesized that this pathway may also be enriched in BCBM and that ACSS2 can regulate tumor growth in the brain parenchyma.

Ferroptosis is a form of non-apoptotic, iron-dependent cell death driven by an excess of lipid peroxides on cellular membranes, leading to membrane damage and regulated non-apoptotic cell death [26]. Accumulation of lipid peroxides and lethal reactive oxygen species (ROS) are controlled by integrated oxidant and antioxidant systems [27]. Cancer cells, and particularly metastasizing ones, undergo high levels of oxidative stress, thus imposing a requirement on their antioxidant system to also be elevated in order to maintain the redox balance. The antioxidant enzyme glutathione peroxidase 4 (GPX4) can convert glutathione (GSH) into oxidized glutathione and directly reduce toxic phospholipid hydroperoxides to nontoxic lipid alcohols, thus acting as a critical repressor of ferroptosis in cancer cells [27]. The amino acid cysteine is considered the main rate-limiting factor in the *de novo* synthesis of GSH. In the high oxidative stress environment of cancer, de novo synthesis of cysteine represents a metabolic demand that cannot be sustained, thus leading to its requirement to be met *via* transmembrane nutrient transporters. In the highly oxidative extracellular space, cysteine exists mainly in its oxidized dimer form, cystine, and it is imported intracellularly through the cystine transporter solute carrier family 7 member 11 (SLC7A11; also, xCT), which is the functional light chain subunit of the system x_c_ ^-^, an L-cystine/L-glutamate antiporter on the cell surface. SLC7A11 exchanges cystine and glutamate across the plasma membrane and cystine is then rapidly reduced into cysteine intracellularly and used to generate GSH [27]. Inhibition of xCT by drugs like erastin can trigger ferroptosis in cancer cells [26]. Whether BCBM cells are susceptible to ferroptosis is not known.

Here, we present evidence that levels of OGT, CDK5 and ACSS2 are elevated in breast cancer brain metastatic (BCBM) trophic cells compared to primary tumor parental breast cancer cells, and their overexpression provides an advantageous growth of primary tumor breast cancer cells in the brain. Importantly, ACSS2 and the phosphomimetic version of ACSS2 (ACSS2-S267D) are required for breast cancer growth only in the brain parenchyma and not the mammary fat pad, contributing to cell survival by protecting against ferroptosis in BCBM cells in an E2F1-dependent manner. In addition, p-Ser267-ACSS2 is highly expressed in patients with brain metastases, providing further evidence for its potential role in the adaptation of breast cancer cells to the brain environment. Importantly, we show for the first time that a novel brain permeable ACSS2 inhibitor can induce ferroptosis and block tumor growth in BCBM cells *ex vivo* and *in vivo*.

## RESULTS

### OGT/CDK5/ACSS2 levels are elevated in breast cancer brain metastatic cells and are required for tumor growth *in vivo*

Given the metabolic similarities in acetate dependency between glioblastoma and brain metastases [13], we hypothesized that our previously identified pathway in GBM [16], where OGT governs regulation of acetate metabolism and growth *via* CDK5 phosphorylation of ACSS2, may also be contributing to BCBM growth. To test this hypothesis, we examined this pathway utilizing human and mammalian parental (PA) and brain trophic (BR) TNBC cells that have been selected to preferentially metastasize to the brain [28]. Initially, to test that these brain tropic cells preferentially grow and have adapted to the brain microenvironment, we intracranially injected parental MDA-MB-231 TNBC cells or the brain tropic breast cancer MDA-MB-231-BR cells enriched to metastasize to the brain, in the parenchyma of immunocompromised Nu/Nu mice [29]. The brain trophic cells formed large and highly invasive tumors compared to parental MDA-MB-231 cells which exhibited smaller, localized focal tumors, indicating a clear adaptation of the BR cells to the brain environment (Supplementary Fig. 1A). Consistent with our hypothesis, we found that the brain trophic cells contained elevated levels of O-GlcNAc and ACSS2-S267 phosphorylation by immunohistochemical (IHC) analysis (Supplementary Fig. 1A) and total levels of O-GlcNAc, OGT and ACSS2 using immunoblot analysis (**Fig. 1A**) when compared to parental breast cancer cells. We saw similar results in two additional pairs of parental and brain trophic mouse TNBC cells: 4T1-BR and E0771-BR cells compared to parental cells (**Fig. 1A**). Importantly, using a tissue microarray, we found that 62% and 94% of human brain cancer metastasis from 86 different primary tumors including breast, lung, thyroid and others contained medium-high levels of O-GlcNAc and ACSS2-S267 phosphorylation, respectively, as determined by immunohistochemical (IHC) analysis (Supplementary Fig. 1B). These findings demonstrate the importance of O-GlcNAc and pS267-ACSS2 in brain metastatic growth.

**Figure 1.**
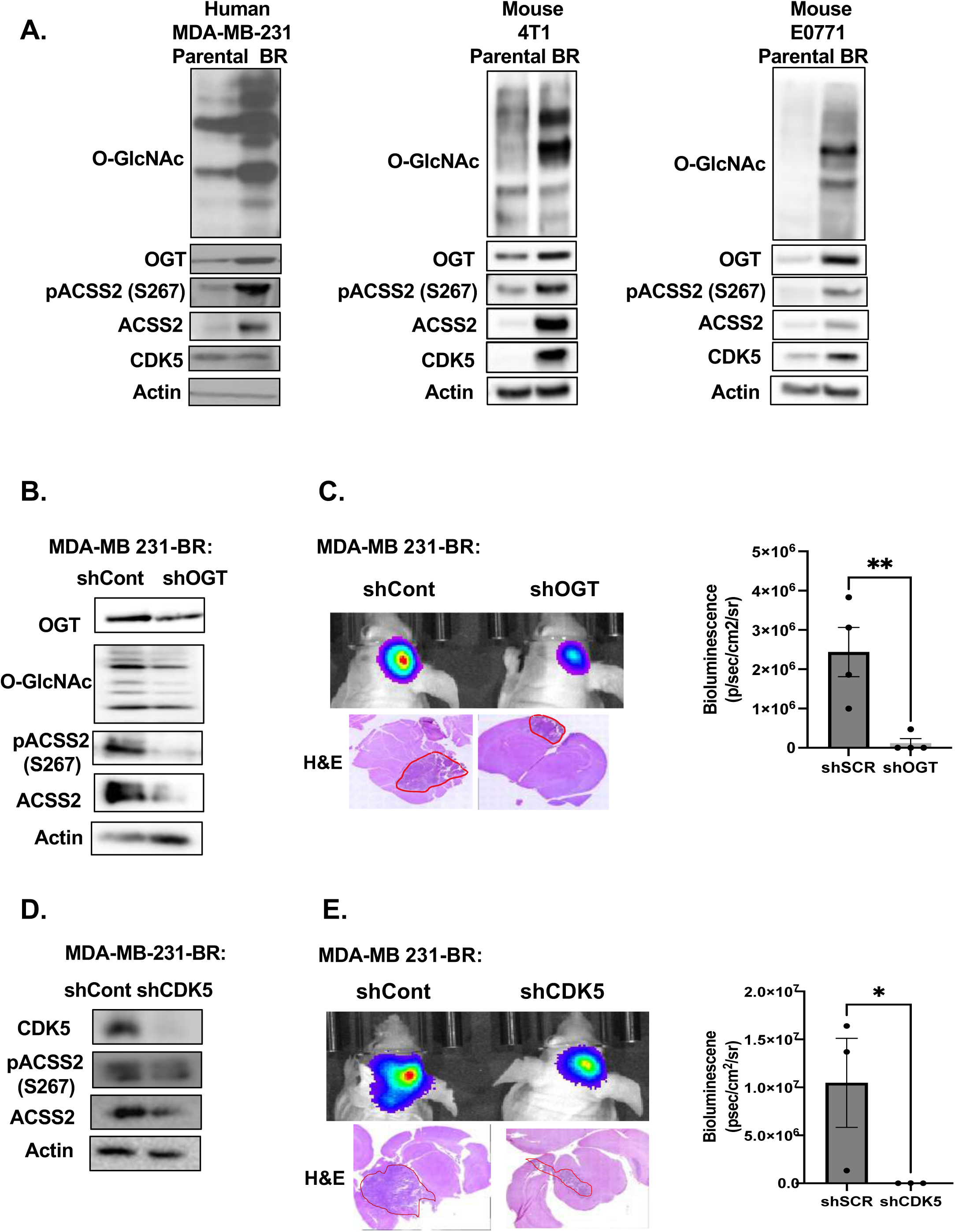
OGT/CDK5/ACSS2 axis is elevated in TNBC BCBM and OGT/CDK5 are required for BCBM cell growth *in vivo*. **(A)** Cell lysate from parental and brain trophic MDA-MB-231, 4T1 and E0771 cells were collected for immunoblot analysis with the indicated antibodies. **(B)** Cell lysates from MDA-MB-231Br cells expressing shRNA against Scramble or OGT were collected for immunoblot analysis with the indicated antibodies. **(C)** Representative images of bioluminescent (top) detection of tumors from mice injected with shSCR and shOGT MDA-MB-231BR cells 14 days post-injection. Representative images of H&E analysis (bottom) on coronal sections from mice harboring shRNA against Scramble or OGT tumors at Day 14. Data are quantified and presented as average Bioluminescence signal from mice injected with MDA-MB-231BR cells expressing shRNA against Scramble (n=3) or OGT mice (n=3) (bottom). Student’s t-test reported as mean ± SEM; **p<0.001. **(D)** Cell lysates from MDA-MB-231Br cells with shRNA against Scramble or CDK5 were collected for immunoblot analysis with the indicated antibodies. **(E)** Representative images of bioluminescent (top) detection of tumors from mice injected with shScramble and shCDK5 MDA-MB-231BR cells 14 days post-injection. Representative images of H&E analysis (bottom) on coronal sections from mice harboring shRNA against Scramble or CDK5 tumors at Day 14. Data are quantified and presented as average Bioluminescence signal from mice injected with MDA-MB-231BR cells expressing shSCR (n=3) or shCDK5 mice (n=3) (bottom). Student’s t-test reported as mean ± SEM; *p<0.05.

To test the role of OGT and CDK5-ACSS2 axis in regulating BCBM macrometastatic growth in the brain, we targeted OGT and CDK5 using stable RNAi knockdown in MDA-MB-231-BR cells injected intracranially. We utilized intracranial injections of breast cancer brain-trophic cells as this is regarded as a suitable model for breast cancer brain macrometastases [30]. Targeting OGT levels in MDA-MB-231-BR cells reduced total O-GlcNAc levels, ACSS2-S267 phosphorylation and total ACSS2 levels compared to control cells (**Fig. 1B**). Intracranial injection of BR cells stably expressing OGT RNAi significantly reduced growth in the brain compared to control cells (**Fig. 1C**). Targeting CDK5 with RNAi in MDA-MB-231-BR cells reduced ACSS2-S267 phosphorylation and total ACSS2 levels compared to control cells (**Fig. 1D**). Similar to the results with reduced OGT expression, intracranial injection of cells stably expressing CDK5 RNAi significantly reduced growth in the brain compared to control cells (**Fig. 1E**). To test whether OGT or CDK5 expression is sufficient to provide signals that allow poorly growing parental cells to adapt to the brain microenvironment and increase growth in the brain parenchyma, we tested the effects of stably overexpressing either OGT or CDK5 in parental MD-MBA-231 cells. OGT overexpressing cells increased O-GlcNAc, CDK5 levels, ACSS2-S267 phosphorylation and total ACSS2 levels compared to controls (Supplementary Fig. 1C) and allowed poorly growing parental breast cancer cells to now form significantly larger tumors in the brain parenchyma compared to control parental MDA-MB-231 cells (Supplementary Fig. 1D). Similar results were obtained in MDA-MB-231 cells overexpressing CDK5 (Supplementary Fig. 1E**, 1F**). These data suggests that OGT and CDK5-mediated phosphorylation of ACSS2 may regulate breast cancer cells metabolic adaptability and growth in the brain microenvironment.

### ACSS2 and ACSS2-pSer267 provide a specific advantage for breast cancer brain metastatic growth

It was previously shown that ACSS2 expression is increased in brain metastases, where similarly to GBM, acetate is the main metabolite to be oxidized in patient tumors [31] [13] [32]. Considering this evidence, we next examined whether ACSS2 and ACSS2-pSer267 are critical for allowing breast cancer cells to adapt to the brain microenvironment. We implanted control and ACSS2 stable RNAi-expressing TNBC cells into the brain or mammary fat pad and assessed tumor growth. Brain tropic MDA-MB-231-BR cells stably expressing ACSS2 RNAi (**Fig. 2A**) exhibited a significantly impaired tumor growth in the brain compared to control cells (**Fig. 2B**). A similar inhibition of tumor growth was seen in brain tropic mouse 4T1-BR cells with stable ACSS2 knockdown compared to controls (Supplementary Fig. 2A, 2B). However, targeting ACSS2 in parental breast cancer MDA-MB-231 cells had minimal effect on tumor growth compared to control when implanted in the mammary fat pad (**Fig. 2C, 2D**). These data suggest a specific dependency on ACSS2 for breast cancer growth in the brain. To directly test the role of ACSS2-S267 phosphorylation on breast cancer growth in the brain, we targeted endogenous ACSS2 using a 3’UTR specific ACSS2 RNAi in parental MDA-MB-231 cells and exogenously expressed either wildtype or ACSS2 phospho-mimetic S267D mutant (**Fig. 2E**). Importantly, growth of tumors derived from ACSS2-S267D expressing cells implanted in the brain was significantly enhanced compared to wildtype ACSS2 expressing cells (**Fig. 2F**), while no significant difference in tumor growth was measured between these same cells when implanted into the mammary fat pad (**Fig. 2G**). These data suggests that ACSS2 and ACSS2-S267 phosphorylation play a unique and critical role in breast cancer cell growth specifically in the brain microenvironment.

**Figure 2.**
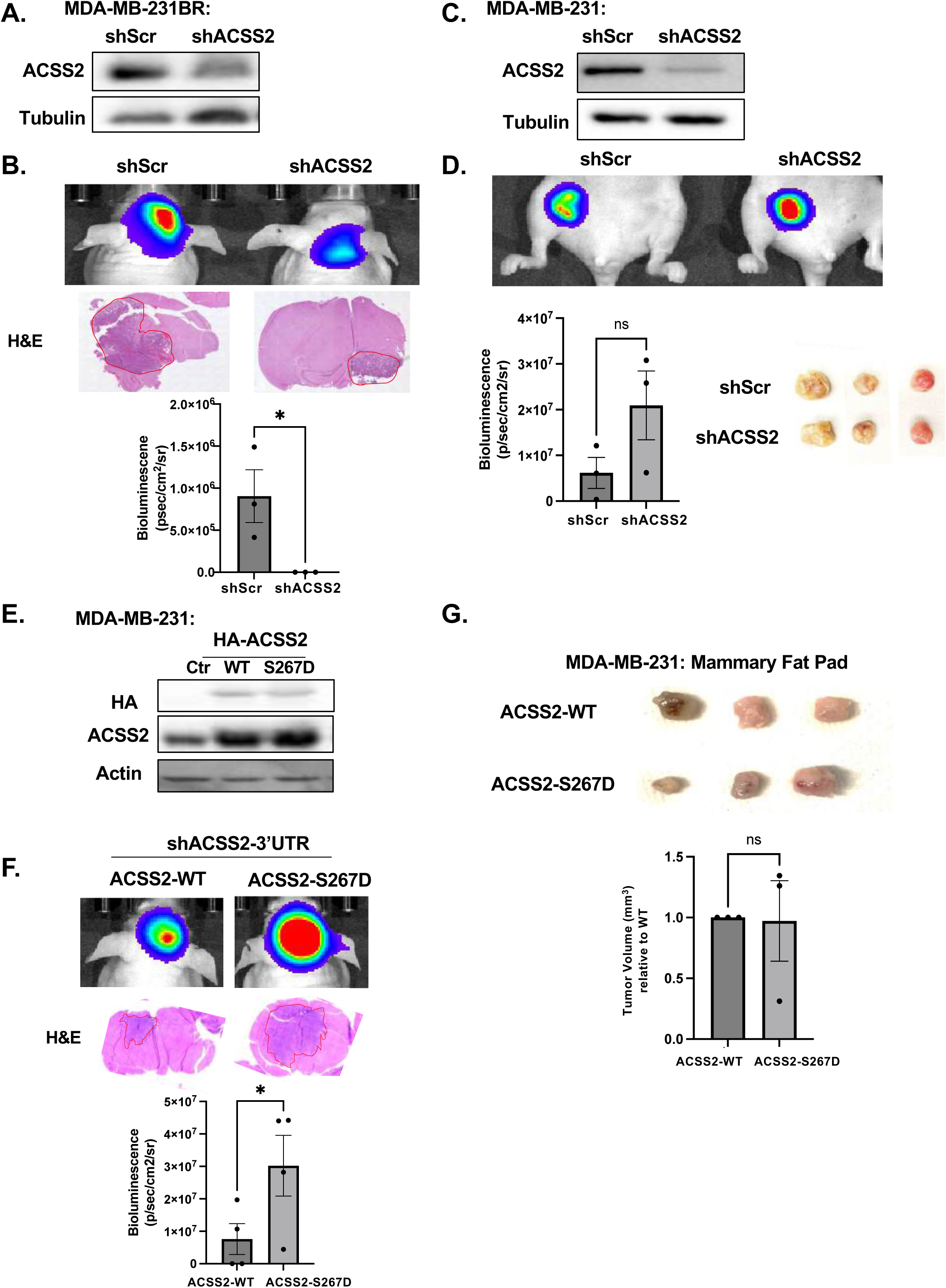
ACSS2 and pSer267-ACSS2 provide a specific growth advantage for tumor growth in the brain but not in the mammary fat pad. **(A)** Cell lysates from MDA-MB-231BR cells stably expressing shRNA against Scramble or ACSS2 were collected for immunoblot analysis with the indicated antibodies. **(B)** Representative images of bioluminescent (top) detection of tumors from mice injected with shSCR and shACSS2 MDA-MB-231BR cells 14 days post-injection. Representative images of H&E analysis (bottom) on coronal sections from mice harboring shRNA against Scramble or ACSS2 tumors at Day 14. Bioluminescence signal from mice injected with MDA-MB-231BR expressing shRNA against Scramble (n=3) or ACSS2 mice (n=3) (bottom). Student’s t-test reported as mean ± SEM; *p<0.05**. (C)** Cell lysates from MDA-MB-231 cells stably expressing shRNA against Scramble or ACSS2 shRNA were collected for immunoblot analysis with the indicated antibodies. **(D)** Representative images of bioluminescent (top) detection of mammary fat tumors from mice injected with shSCR and shACSS2 MDA-MB-231 cells 21 days post-injection. Representative images of fat pad tumors removed (bottom) from mice harboring shRNA against Scramble or ACSS2 tumors at Day 21. Bioluminescence signal from mice injected with MDA-MB-231 expressing shRNA against Scramble (n=3) or ACSS2 mice (n=3) (bottom). Student’s t-test reported as mean ± SEM; n.s.. **(E)** Cell lysates from MDA-MB-231 cells stably expressing shRNA against endogenous ACSS2, overexpressing HA-tagged wildtype (WT) or HA-tagged phosphomimetic mutant (S267D) ACSS2 were collected for immunoblot analysis with the indicated antibodies. **(F)** Representative images of bioluminescent (top) detection in MDA-MB-231 cells with shRNA against endogenous ACSS2, overexpressing HA-tagged wildtype (WT) or HA-tagged phosphomimetic mutant (S267D). Representative images of H&E analysis (bottom) on coronal sections from mice harboring shRNA against endogenous ACSS2, overexpressing HA-tagged wildtype (WT) or HA-tagged phosphomimetic mutant (S267D) ACSS2 tumors at Day 14. Bioluminescence signal from mice injected with MDA-MB-231 expressing shRNA against Scramble, overexpressing HA-tagged wildtype (WT) (n=4) or HA-tagged phosphomimetic mutant (S267D) ACSS2 mice (n=4) (bottom). Student’s t-test reported as mean ± SEM; *p<0.05. **(G)** Representative images of fat pad tumors removed (bottom) from mice harboring shRNA against endogenous ACSS2, overexpressing HA-tagged wildtype (WT) or HA-tagged phosphomimetic mutant (S267D) ACSS2 tumors at Day 21. Student’s t-test reported as mean ± SEM; n.s..

We also tested whether the role of CDK5 in regulating BCBM growth required ACSS2-S267 phosphorylation. To test whether CDK5-mediated regulation of cell growth was dependent on ACSS2-S267 phosphorylation, we overexpressed ACSS2-S267D mutant in the context of CDK5 knockdown (Supplementary Fig. 3A) and noticed a partial growth restoration in MDA-MB-231BR cells *in vitro* (Supplementary Fig. 3B). In addition, MDA-MB-231-BR cells expressing the phospho-mimetic mutant ACSS2-S267D were able to partly rescue the growth effects conferred by CDK5 depletion *in vivo* (Supplementary Fig. 3C). Thus, CDK5 regulation of BCBM growth is partly dependent on ACSS2-S267 phosphorylation.

### ACSS2 regulates breast cancer brain metastatic cell survival via ferroptosis suppression

Using an *ex vivo* brain slice tumor model [33], we treated brain slices containing MDA-MB-231-BR preformed tumors with a quinoxaline-based ACSS2 inhibitor, VY-3-135 [34] and a previous chemotype VY-3-249 (ACSS2i) (**Fig. 3A)** and found that targeting ACSS2 causes a complete regression of tumors in the brain parenchyma (**Fig. 3B**) compared to control treatment, suggesting that targeting ACSS2 induced cell death in BCBM cells. To exclude toxic traits of the VY-3-249 ACSS2 inhibitor in *ex vivo* brain slices, we examined brain slice viability in tumor-free slices and found no difference with vehicle treated samples, indicating that the effect of VY-3-249 is specific to the cancer cells (Supplementary Fig. 4A). A four-fold increase in propidium iodide (PI) in MDA-MB-231-BR cells treated with VY-3-249 confirmed induction of cell death (**Fig. 3C**). To determine the specific cell death mechanism mediated by targeting ACSS2, we tested several cell death inducing inhibitors including autophagy (3-methyladenine: 3-MA), necroptosis (necrosulfonamide: NSA), apoptosis (caspase inhibitor: ZVAD) and ferroptosis (ferrostatin-1: Fer-1). Only Ferrostatin-1 (Fer-1), a specific inhibitor of ferroptosis [35], blocked VY-3-249 induced cell death (**Fig. 3C**). Ferroptosis is an iron-dependent cell death mechanism characterized by the accumulation of lipid-peroxides that leads to plasma membrane rupture [36]. We detected a near seven-fold increase in lipid peroxides in MDA-MB-231-BR cells treated with VY-3-249, as measured by C11 BODIPY 581/591 staining (**Fig. 3D**). This effect was comparable to treating cells with erastin (**Fig. 3D**), an inhibitor of system x_c_ ^-^ antiporter (xCT), and a well-characterized drug that triggers ferroptosis [37]. We detected a similar increase in cell death (Supplementary Fig. 4B) and lipid peroxidation (Supplementary Fig. 4C) in 4T1-BR cells treated with VY-3-249. In both cell lines, these effects were reversed by treatment with Fer-1. Consistent with these data, targeting ACSS2 with RNAi in MDA-MB-231-BR (**Fig. 3E**) and 4T1-BR cells (Supplementary Fig. 4D) led to significant increase in cell death (**Fig. 3F, Supplementary** Fig. 4E) and lipid peroxides (**Fig. 3G, Supplementary** Fig. 4F) which were reversed by treatment with Fer-1.

**Figure 3.**
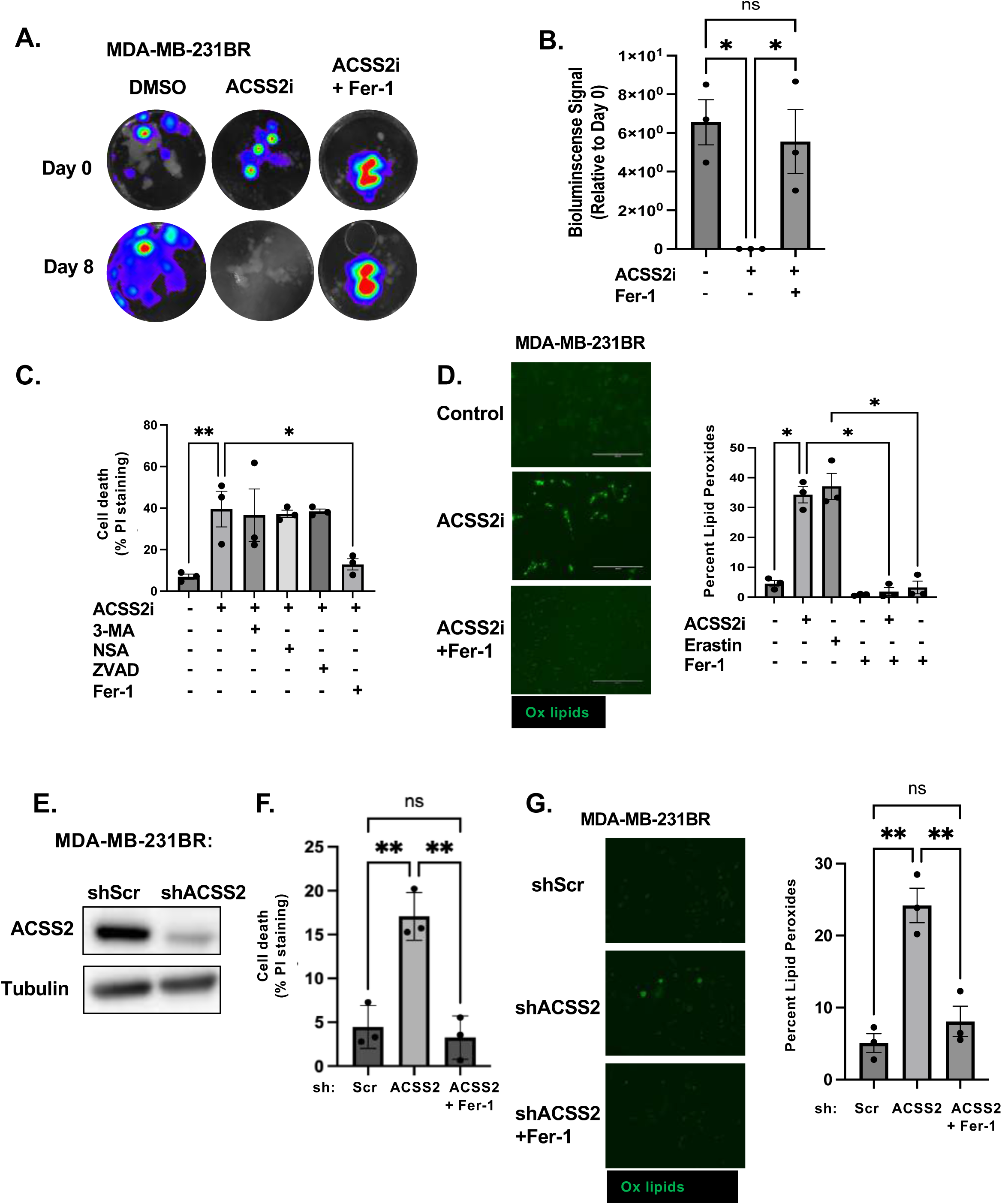
ACSS2 regulates breast cancer brain metastatic cell survival via Ferroptosis suppression. **(A)** Representative images depicting tumor growth in organotypic brain slices derived from mice intracranially injected with MDA-MB-231BR-luc cells detected via bioluminescence. Brain slices containing tumors are treated with Control (DMSO), ACSS2 inhibitor (ACSS2i) 50 μM or ACSS2i and Ferrostatin-1 (Fer-1) 5 μM for indicated days (top). **(B)** Quantification of tumor Bioluminescence at indicated day (Control: DMSO n=3, ACSS2i n=3, ACSS2i +Fer-1 n=3) One-way ANOVA with Tukey’s multiple comparisons test reported as mean ± SEM *p-value < 0.05. **(C)** Quantification of propidium iodine staining flow cytometry of MDA-MB-231BR cells treated with ACSS2 inhibitor (ACSS2i) (n=3), ACSS2i + 3-Methyladenine (3-MA) (n=3), ACSS2i + Necrosulfonamide (NSA) (n=3), ACSS2i + Caspase Inhibitor Z-VAD-FMK (ZVAD) (n=3), or ACSS2i + Ferrostatin-1 (Fer-1). Data reported as One-way ANOVA with Tukey’s multiple comparisons test reported as mean ± SEM *p-value < 0.05, **p<0.01. **(D)** Representative images of MDA-MB-231BR cells treated with Control (DMSO), ACSS2i, or ACSS2i + Fer-1 and stained with Bopidy C11 (image magnification 20x, scale bar: 200 µM) (right). Quantification of lipid peroxides with Bopidy C11 and flow cytometry of MDA-MB-231BR cells treated with ACSS2i, Erastin, Fer-1, ACSS2i + Fer1, Erastin + Fer-1. Data reported as One-way ANOVA with Tukey’s multiple comparisons test reported as mean ± SEM *p-value < 0.05. **(E)** Cell lysates from MDA-MB-231BR cells stably expressing shRNA against Scramble or ACSS2 were collected for immunoblot analysis with the indicated antibodies. **(F)** Quantification of propidium iodine staining flow cytometry of MDA-MB-231BR cells expressing shRNA against Scramble, ACSS2, and shRNA against ACSS2 and Fer-1. Data reported as One-way ANOVA with Tukey’s multiple comparisons test reported as mean ± SEM **p-value < 0.01. **(G)** Representative images of MDA-MB-231BR cells stably expressing shRNA against Scramble or ACSS2 and treated with + Fer-1 and stained with Bopidy C11 (image magnification 20x, scale bar: 200 µM) (right). Quantification of lipid peroxides with Bopidy C11 and flow cytometry of MDA-MB-231BR cells expressing shRNA against Scramble, ACSS2, and shRNA against ACSS2 and Fer-1 (left). Data reported as One-way ANOVA with Tukey’s multiple comparisons test reported as mean ± SEM **p-value < 0.01.

We next examined whether Fer-1 treatment could also block tumor regression following VY-3-249 treatment in the *ex vivo* tumor brain slice model. First, we show that preformed tumors exhibited significantly reduced tumor growth when treated with the ferroptosis inducer erastin, while cotreatment with Fer-1 blocked this effect (Supplementary Fig. 4G), indicating that *ex vivo* brain slices can undergo ferroptotic regulation. Consistent with *in vitro* results (**Fig. 3C**), VY-3-249 mediated effects on tumor growth *ex vivo* were reversed by Fer-1 co-treatment (**Fig. 3A, 3B**). Thus, targeting ACSS2 genetically and pharmacologically induces ferroptosis in BCBM cells *in vitro* and *ex vivo*.

### ACSS2 regulates ferroptosis in an E2F1-dependent manner in BCBM cells

To explore downstream mechanisms by which ACSS2 regulates ferroptosis in BCBM cells, we performed bulk RNA-sequencing (RNAseq) analysis to identify transcriptional changes in gene expression patterns in MDA-MB-231-BR cells containing downregulated levels of ACSS2 by RNAi. Transcriptome analysis of MDA-MB-231-BR cells comparing stable expression of ACSS2 RNAi to non-gene targeting RNAi identified 422 differentially expressed genes consisting of 168 upregulated and 254 downregulated genes using a cut-off value of fold change >2 (and p<0.05); (Supplementary Fig. 5A, **Supplementary Table S1**). Our RNAseq analysis did not contain changes in major ferroptosis regulators. However, expression of transcription factor E2F1-regulated genes was significantly decreased upon ACSS2 knockdown (Supplementary Fig. 5B). E2F1 is a central player involved in cell cycle and cell death and has recently been implicated in the regulation of ferroptosis [38–40]. Consistent with the RNA-seq analysis, we found that knockdown of ACSS2 leads to downregulation of E2F1 protein (**Fig. 4A**) and RNA levels (**Fig. 4B**) in MDA-MB-231BR cells and E2F1 protein levels in 4T1-BR cells (Supplementary Fig. 4D). Consistent with our hypothesis that ACSS2 regulation of E2F1 may be linked to ferroptosis, we found that targeting E2F1 with RNAi in MDA-MB-231BR cells (**Fig. 4C**) significantly increases cell death (**Fig. 4D**) and lipid peroxide levels (**Fig. 4E, 4F**) that were reversed by Fer-1 co-treatment (**Fig. 4D, 4E, 4F**). Conversely, we show that overexpression of E2F1 in cells containing RNAi depleted levels of ACSS2 (**Fig. 4G**) can partly reverse ACSS2-mediated induction of cell death (**Fig. 4H**) and fully rescue lipid peroxide levels (**Fig. 4I**) compared to controls. In addition, we show that E2F1 overexpression can block cell death-mediated effects of treatment with ACSS2 inhibitor (Supplementary Fig. 5C).

**Figure 4.**
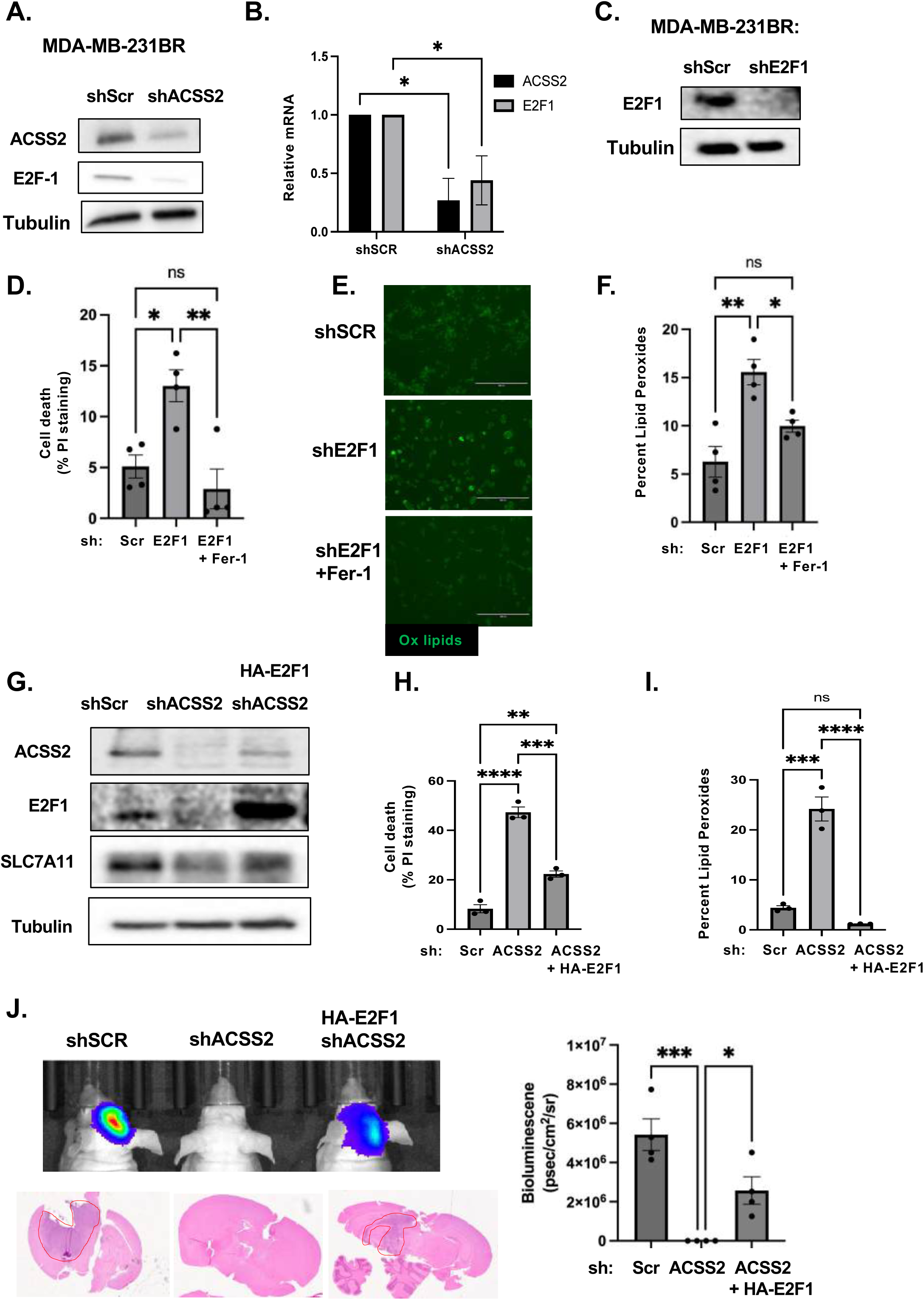
ACSS2 regulates Ferroptosis in an E2F1-dependent regulation of SLC7A11. **(A)** Cell lysates from MDA-MB-231BR cells stably expressing shRNA against Scramble or ACSS2 were collected for immunoblot analysis with the indicated antibodies. **(B)** Total RNA was collected from MDA-MB-231BR cells stably expressing shRNA against Scramble or ACSS2. Quantification of qRT-PCR performed on RNA extracts analyzing ACSS2 and E2F1 gene expression normalized to PPIA. Two-way ANOVA reported as mean ± SEM.* p-value < 0.05. **(C)** Cell lysates from MDA-MB-231BR cells stably expressing shRNA against Scramble or E2F1 were collected for immunoblot analysis with the indicated antibodies. **(D)** Quantification of propidium iodine staining flow cytometry of MDA-MB-231BR cells expressing shRNA against Scramble or E2F1, treated with and without Fer-1. Data reported as One-way ANOVA with Tukey’s multiple comparisons test reported as mean ± SEM *p-value < 0.05. **(E)** Representative images of MDA-MB-231BR cells expressing shRNA against Scramble or E2F1, and Fer-1 Bopidy C11 (image magnification 20x, scale bar: 200 µM) (left). (**F**) Quantification of lipid peroxides with Bopidy C11 and flow cytometry (right) of MDA-MB-231BR cells expressing shRNA against Scramble or shRNA against E2F1 and Fer-1. Data reported as One-way ANOVA with Tukey’s multiple comparisons test reported as mean ± SEM *p-value < 0.05. **(G)** Cell lysates from MDA-MB-231BR cells stably expressing shRNA against Scramble or shRNA against ACSS2 and overpression of HA-Tagged E2F1 were collected for immunoblot analysis with the indicated antibodies. **(H)** Quantification of propidium iodine staining flow cytometry of MDA-MB-231BR cells expressing shRNA against Scramble, shRNA against ACSS2 and HA-tagged E2F1. Data reported as One-way ANOVA with Tukey’s multiple comparisons test reported as mean ± SEM **p-value < 0.01, ***p<0.001, ****p<0.0001. **(I)** Quantification of lipid peroxides with Bopidy C11 and flow cytometry of MDA-MB-231BR cells expressing shRNA against Scramble, ACSS2, or shRNA against ACSS2 and HA-Tagged E2F1 RNA. Data reported as One-way ANOVA with Tukey’s multiple comparisons test reported as mean ± SEM ***p-value < 0.001, ****p<0.0001. **(J)** Representative images of bioluminescent (top) detection of tumors from mice injected with shSCR and shACSS2, shACSS2 HA-E2F1 MDA-MB-231BR cells 14 days post-injection. Representative images of H&E analysis (bottom) on coronal sections from mice as above. Bioluminescence signal was quantified from mice injected with MDA-MB-231BR cells stably expressing shSCR (n=4) and shACSS2 (n=4), shACSS2 HA-E2F1 (n=4) (right). Student’s t-test reported as mean ± SEM; *p<0.05.

Importantly, we show that overexpression of E2F1 in cells containing RNAi targeted ACSS2 levels can partly reverse tumor growth *in vivo* (**Fig. 4J**). E2F1 protein levels are elevated in MDA-MB-231-BR (Supplementary Fig. 5D), 4T1-BR (Supplementary Fig. 5E), and E0771-BR cells (Supplementary Fig. 5F) compared to corresponding parental cells, which is consistent with our data that describe elevated ACSS2 levels in three different breast cancer brain topic lines and further supports ACSS2 dependent regulation of E2F1 expression. Therefore, these data suggest that targeting ACSS2 genetically or pharmacologically induces ferroptotic cell death in an E2F1-dependent manner.

To understand the mechanism by which ACSS2-E2F1 axis regulates ferroptosis, we examined the expression of several key anti-ferroptotic regulators including SLC7A11 and GPX4 which control glutathione metabolism and reduce levels of lipid peroxides [27]. We found that targeting E2F1 genetically reduces the protein (**Fig. 5A**) and RNA (**Fig. 5B**) levels of SLC7A11 and GPX4 in MDA-MB-231BR cells as well as protein levels in 4T1-BR cells (**Fig. 5C**). Conversely, we detected an increase in SLC7A11 and GPX4 protein levels in MDA-MB-231 cells overexpressing E2F1 (Supplementary Fig. 6A). We found that genetically targeting ACSS2 with RNAi reduces RNA (Supplementary Fig. 6B) and protein levels of SLC7A11 and GPX4 in MDA-MB-231-BR (**Fig. 5D**), and protein levels in 4T1-BR (**Fig. 5E**) and E0771-BR (**Fig. 5F**) cells. In addition, the reduction of SLC7A11 and GPX4 levels in MDA-MB-231-BR in ACSS2 RNAi targeted cells was reversed by overexpression of E2F1 (**Fig. 4G**). Further, pharmacologically targeting ACSS2 with VY-3-249 treatment reduced expression of E2F1, SLC7A11 and GPX4 in MDA-MB-231BR (Supplementary Fig. 6C) and 4T1-BR (Supplementary Fig. 6D) cells. All these results suggest that ACSS2 regulates ferroptotic factors SLC7A11 and GPX4 *via* E2F1.

**Figure 5.**
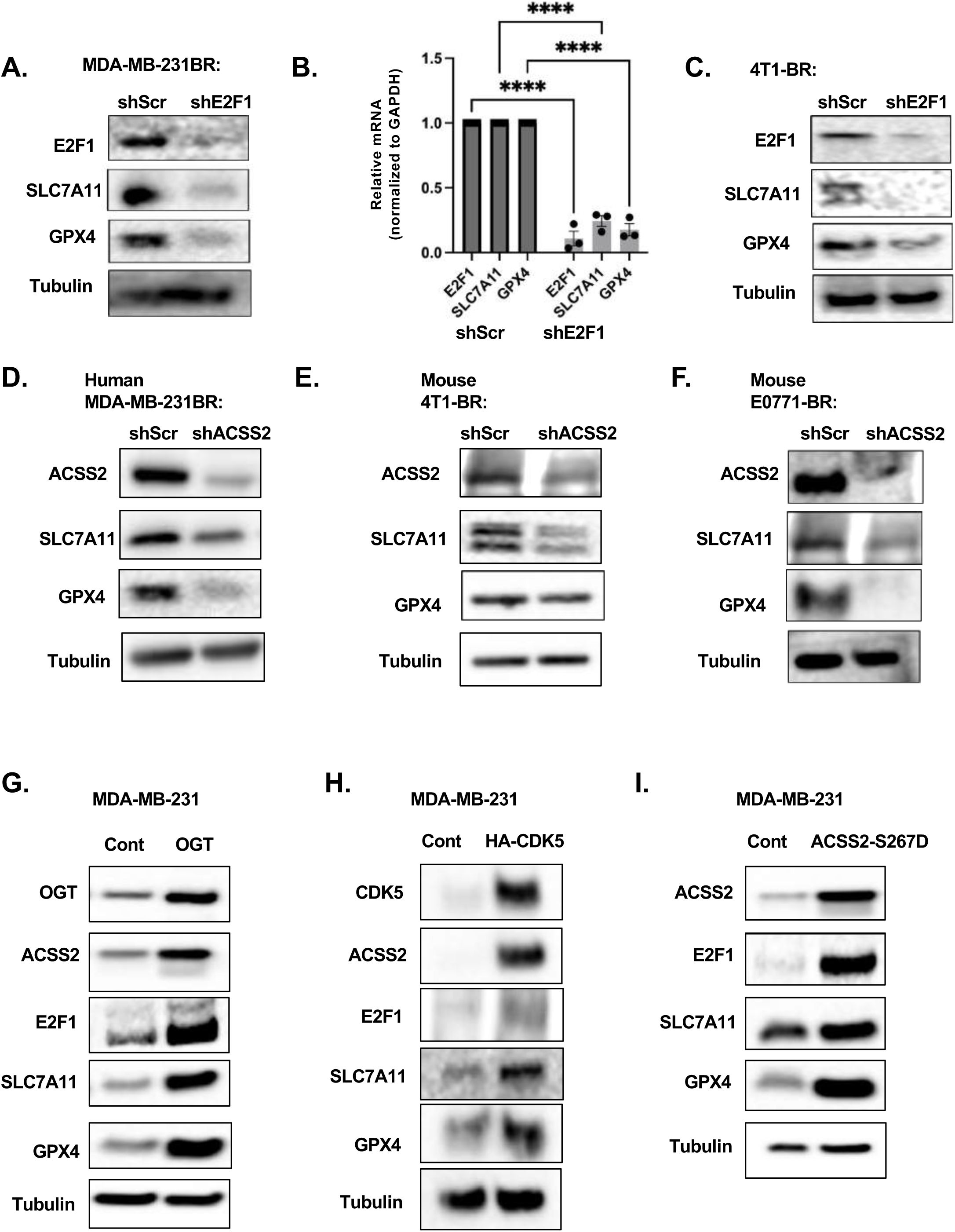
OGT/CDK5/ACSS2 Signaling axis regulates ferroptosis regulators. **(A)** Cell lysates from MDA-MB-231BR cells stably expressing shRNA against Scramble or E2F1 were collected for immunoblot analysis with the indicated antibodies. **(B)** Total RNA was collected from MDA-MB-231BR cells stably expressing shRNA against Scramble or E2F1. Quantification of qRT-PCR performed on RNA extracts analyzing E2F1, SLC7A11, and GPX4 gene expression normalized to GAPDH. Two-way ANOVA reported as mean ± SEM.* p-value < 0.05. **(C)** Cell lysates from 4T1BR cells stably expressing shRNA against Scramble or E2F1 were collected for immunoblot analysis with the indicated antibodies. **(D)** Cell lysates from MDA-MB-231BR cells stably expressing shRNA against Scramble or ACSS2 were collected for immunoblot analysis with the indicated antibodies**. (E)** Cell lysates from 4T1-BR cells stably expressing shRNA against Scramble or ACSS2 were collected for immunoblot analysis with the indicated antibodies. **(F)** Cell lysates from E0771-BR cells stably expressing shRNA against Scramble or ACSS2 were collected for immunoblot analysis with the indicated antibodies. **(G)** Cell lysates from MDA-MB-231 cells stably overexpressing Control or OGT were collected for immunoblot analysis with the indicated antibodies**. (H)** Cell lysates from MDA-MB-231 cells stably overexpressing Control or CDK5 were collected for immunoblot analysis with the indicated antibodies. **(I)** Cell lysates from MDA-MB-231 cells stably overexpressing Control or HA-ACSS2-S267D were collected for immunoblot analysis with the indicated antibodies.

To link these findings to the previously established pathway driven by OGT/CDK5/p-Ser267-ACSS2 upregulation in BCBM, that drives large tumor formation in the brain parenchyma, we show that MDA-MB-231 parental cells overexpressing OGT (**Fig. 5G),** CDK5 (**Fig. 5H**) and ACSS2-S267D mutant (**Fig. 5I**) contained increased protein levels of E2F1, SLC7A11 and GPX4, suggesting an ACSS2 dependent regulation of E2F1, which in turn drives the regulation of anti-ferroptotic proteins SLC7A11 and GPX4.

### Novel brain permeable ACSS2 inhibitor induces ferroptosis and block BCBM growth *ex vivo* and *in vivo*

We have recently identified novel brain-permeable ACSS2 inhibitors, including compound AD-5584, that is predicted to be more stable than both VY-3-249 and VY-3-135 and can cross the blood-brain barrier [34]. Consistent with the data above, treatment of BCBM cells with AD-5584 reduces E2F1, SLC7A11, and GPX4 protein levels (**Fig. 6A**) and significantly increases cell death (**Fig. 6B**) and lipid peroxides (**Fig. 6C**) in MDA-MB-231BR cells compared to control-treated cells, which were reversed by treatment with Fer-1. Similar results are shown for 4T1-BR cells treated with AD-5584 *in vitro* (Supplementary Fig. 7A, B, C. Consistent with the model that ACSS2-mediates ferroptotic resistance *via* regulation of E2F1, we show that overexpression of E2F1 in MDA-MB-231-BR cells protects against AD-5584-mediated cell death (**Fig. 6D**) and lipid peroxidation (**Fig. 6E**). We also found that treatment with novel ACSS2 inhibitor AD-5584 was able to cause tumor regression in preformed tumors *ex vivo,* which was reversed by Fer-1 co-treatment in MDA-MB-231BR (**Fig. 6F**) and 4T1-BR tumors (Supplementary Fig. 7D). Importantly, we show that treating mice with a brain permeable ACSS2 inhibitor can significantly block breast cancer brain metastatic growth of 4T1BR (**Fig. 6G**) and MDA-MB-231BR (Supplementary Fig. 7E) *in vivo* and significantly extend survival of mice compared to control treatment (**Fig. 6H**). This inhibitor did not have any observable toxicity *ex vivo,* as treatment of tumor free brain slices with AD-5584 did not alter brain slice viability [41]. In addition, animals treated with AD-5584 did not lose weight compared to controls (Supplementary Fig. 7F, 7G) or have any observable change in behavior. Thus, we show that targeting BCBM tumors with a novel brain-permeable ACSS2 inhibitor induces ferroptosis *in vitro* and blocks tumor growth *ex vivo* and *in vivo*.

**Figure 6.**
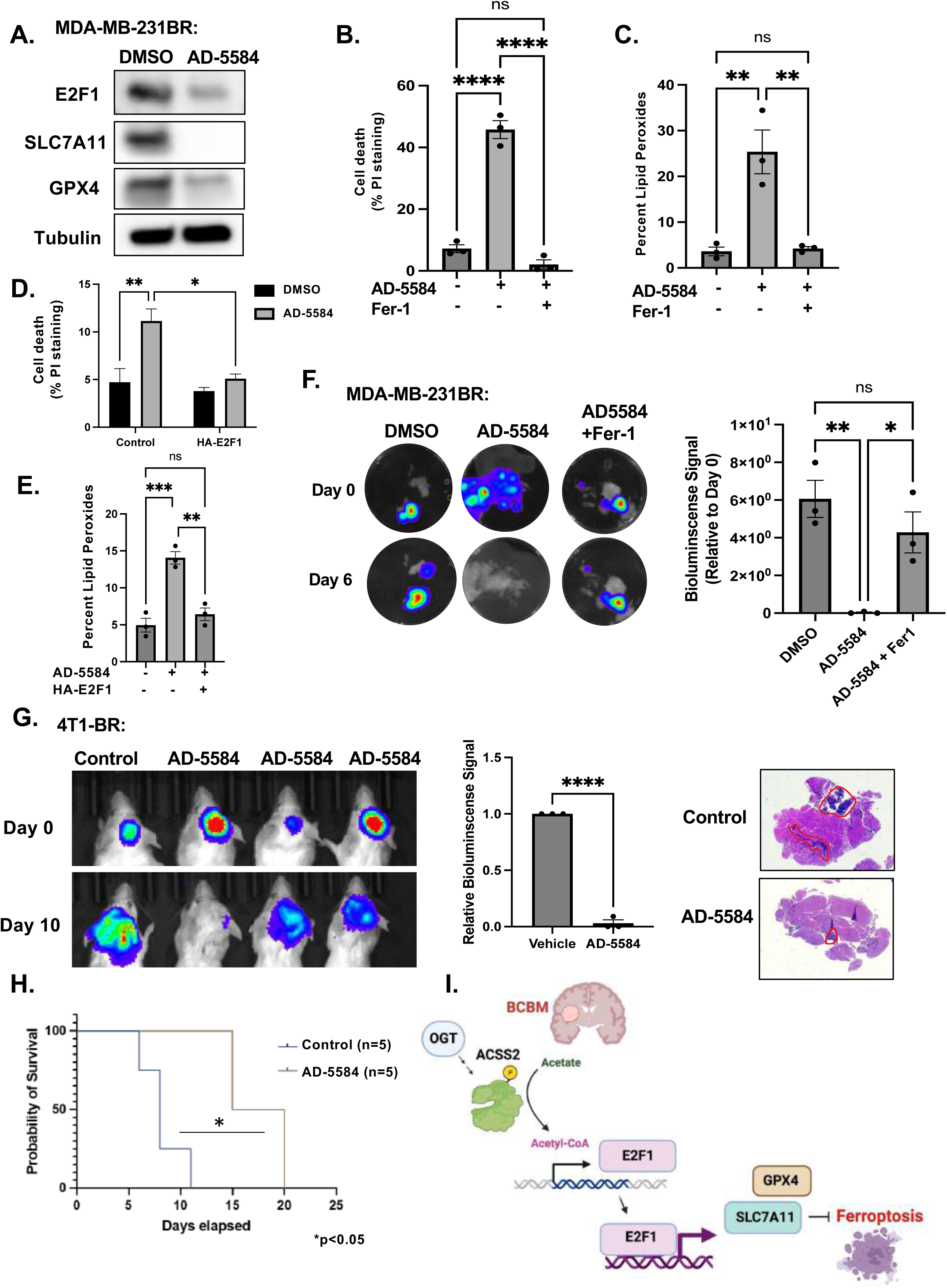
Novel Brain Permeable ACSS2 inhibitor induces Ferroptosis in an E2F1-dependent pathway. **(A)** Cell lysates from MDA-MB-231BR cells treated with control (DMSO) or ACSS2 inhibitor AD-5584 (100 μM) for 48 hrs were collected for immunoblot analysis with the indicated antibodies. (**B)** Quantification of propidium iodine staining flow cytometry of MDA-MB-231BR cells treated with DMSO, AD-5584, or AD-5584 + Fer1. Data reported as One-way ANOVA with Tukey’s multiple comparisons test reported as mean ± SEM *p-value < 0.05, **p<0.01. **(C)** Quantification of lipid peroxides with Bopidy C11 and flow cytometry of MDA-MB-231BR (right) cells treated with DMSO, AD-5584, or AD-5584 + Fer1. Data reported as One-way ANOVA with Tukey’s multiple comparisons test reported as mean ± SEM **p-value < 0.01. **(D)** Quantification of propidium iodine staining flow cytometry of MDA-MB-231BR cells overexpressing control or E2F1 treated with DMSO, AD-5584, or AD-5584. Data reported as One-way ANOVA with Tukey’s multiple comparisons test reported as mean ± SEM *p-value < 0.05, **p<0.01 **(E)** Quantification of lipid peroxides with Bopidy C11 and flow cytometry of MDA-MB-231BR cells overexpressing E2F1 treated with DMSO, AD-5584, or AD-5584. Data reported as One-way ANOVA with Tukey’s multiple comparisons test reported as mean ± SEM **p-value < 0.01, ***p-value < 0.001. **(F)** Representative images depicting tumor growth in organotypic brain slices derived from mice intracranially injected with MDA-MB-231Br-luc cells detected via bioluminescence. Brain slices containing tumors are treated with Control (DMSO), AD-5584 or AD-5584 and Ferrostatin-1 (Fer-1) for indicated days (left). Quantification of tumor Bioluminescence at indicated day (right) (Control: DMSO n=3, AD-5584 n=3, AD-5584 +Fer-1 n=3) One-way ANOVA with Tukey’s multiple comparisons test reported as mean ± SEM *p-value < 0.05, **p<0.01. **(G)** Representative images of bioluminescent detection of tumors from Balb/C mice injected with luciferase-tagged 4T1BR cells at Day 0 (prior to drug treatment) and at 10 days post-drug treatment. Data are quantified and presented as average Relative Bioluminescence signal from mice injected with 4T1BR cells treated with Vehicle (n=3) or AD-5584 treated mice (n=3) (right). Student’s t-test reported as mean ± SEM; *p<0.05. Representative images of brain sections stained for H&E 10-days-post treatment. **(H)** Kaplan-Meyer survival analysis quantifying survival of mice injected with 4T1BR cells and treated with vehicle (n=5) or AD-5584 (n=5), *p<0.05. **(I)** Working model schematic depicting OGT regulates ACSS2 via CDK5 phosphorylation at Serine-267 residue. BCBM cells upregulate ACSS2 to convert acetate to acetyl-CoA. Acetyl-CoA promotes the transcription of E2F1, leading to the transcription of anti-ferroptotic E2F1 target genes, SLC7A11 and GPX4. Expression of SLC7A11and GPX4 protects BCBM cells from ferroptosis.

## DISCUSSION

Colonization and eventual growth in the brain microenvironment by cancer cells requires a unique metabolic adaptation that involves a dependency on acetate as a metabolic fuel and increased ACSS2 levels. In the present study, we have unveiled molecular mechanisms underlying cell growth and survival pathways of breast cancer brain metastatic cells. We have shown that the OGT/CDK5/ACSS2 axis can regulate breast cancer brain metastatic cell growth in the brain parenchyma, in part, via ACSS2 protection from ferroptosis in a E2F1-dependent manner. Interestingly, ACSS2 was critical for growth in breast cancer cells in the brain but not mammary fat pad *in vivo*. This result in the mammary fat pad aligns with recent finding showing that ACSS2 knockdown has no effect in the mammary fat pad of immune-deficient mice [42]. Thus, ACSS2 may represent a critical regulator of breast tumors in the brain parenchyma. Moreover, we showed that ACSS2 protects against ferroptosis in BCBM cells in a E2F1-dependent manner. Our model suggests that ACSS2 may likely regulate acetylation on key transcriptional regulators on the E2F1 promoter, and E2F1, in turn, can regulate expression of SLC7A11 and GPX4 (**Fig. 6H**) to protect against ferroptosis. In neurons, ACSS2 regulates gene expression through direct binding to chromatin to promote histone acetylation [43] and motif analysis of ACSS2-targeted, acetate-induced genes implicates the regulation of transcription factors including E2F3 [44]. Thus, ACSS2 in breast cancer brain metastatic cells may serve to protect from ferroptosis via regulation of gene expression.

ACSS2 has emerged as a critical regulator of tumor growth in a number of cancers [45] by affecting cancer lipid metabolism and autophagy [45]. Our study is the first to link ACSS2 in regulating ferroptosis in brain tropic breast cancer cells. In addition, we found that targeting ACSS2 genetically or pharmacologically increased lipid peroxidation in multiple BCBM cells. We found that ACSS2 regulates E2F1 levels and E2F1-regulated genes in breast cancer brain metastatic trophic cells and that ACSS2 regulation of ferroptosis is E2F1 dependent. E2F1 alone was able to regulate ferroptosis and targeting E2F1 induces cell death and lipid peroxidation in BCBM cells, which were reversed by ferrostatin-1 treatment. These data are consistent with recent studies showing E2F1 as a ferroptosis regulator in cancer cells. In prostate cancer, RB1 loss/E2F activation sensitized cancer cells to ferroptosis [40], while in fibrosarcoma cells, E2F1 overexpression protected cells from ferroptotic inducers, including erastin, and this protection was associated with E2F1 regulation of both GPX4 and SLC7A11 expression [39]. Interestingly, we found brain tropic cells contain elevated levels of E2F1compared to parental breast cancer cells suggesting that E2F1-mediated protection from ferroptosis may be a mechanism for breast cancer cells to adapt to the unique brain microenvironment. Taken together, our data suggest that ACSS2/E2F1 axis can regulate ferroptosis in breast cancer brain tropic cells.

Since the brain microenvironment is deficient in several nutrients required by cancer cells, including free fatty acids, tumors in the brain, including BCBM, must increase tumor fatty acid synthesis [46]. ACSS2 can contribute to de-novo fatty acid synthesis by converting acetate to acetyl-CoA which is used to generate fatty acids [47]. Ferroptosis is highly dependent on lipid metabolism as lipids are peroxidated and cause membrane disruption, leading to outer membrane rupture [48]. Since BCBM cells contain elevated fatty acid synthesis, this may make them sensitive to ferroptosis and require mechanisms to protect them from this form of cell death in the brain microenvironment. Our data suggests that BCBM cells increase ACSS2 levels to protect them from ferroptosis by regulating E2F1-mediated gene expression of anti-ferroptotic proteins SLC7A11 and GPX4. Whether ACSS2 regulation of lipid synthesis can also contribute to regulation of ferroptosis in BCBM cells needs further examination.

Small molecules targeting ACSS2 are in clinical trials for the treatment of advanced tumors and have been found to be well tolerated in Phase 1 clinical trials [49] (NCT04990739), which is consistent with the finding that Acss2-knockout mice are viable [47]. We have previously identified two novel inhibitors that can cross the blood-brain barrier and show efficacy in pre-clinical models of BCBM [41] (**Fig. 6**). Using an *ex vivo* BCBM tumor model, we showed that targeting ACSS2 may also synergize with radiation treatment [41]. Since ferroptosis is a form of inflammatory cell death [50] that can increase the efficacy of immune therapy in breast cancer [51], targeting ACSS2 may alter immune cells in the brain tumor microenvironment and sensitize them to immune checkpoint inhibitors. Indeed, recent studies have shown that targeting ACSS2 may also regulate the host immune response to cancer [42]. Human genomic analysis found that ferroptotic markers are associated with poor clinical outcomes in BCBM patients [52]. Thus, targeting ACSS2 in breast cancer brain metastatic cells may have clinical utility in inducing cell death via ferroptosis without the toxicity seen with agents such as erastin [53] and may synergize in combination with cancer therapies including radiotherapy or immunotherapy. In summary, our study provides a framework for targeting BCBM tumors with brain-penetrant ACSS2 inhibitors to induce ferroptotic cell death to exploit the metabolic vulnerability of these tumors.

## METHODS

### Cell Culture

Triple negative breast cancer MDA-MB-231, 4T1, HEK293T cell lines were obtained from ATCC (Manassas, VA, USA) and cultured according to ATCC protocol supplemented with 10% fetal bovine serum (FBS), 5% 10,000 Units/mL Penicillin-10,000 μg/mL Streptomycin, and 5% 200 mM L-Glutamine. Triple-negative brain trophic cells MDA-MB-231BR, and 4T1BR are a kind gift from Dr. Patricia Steeg (Center for Cancer Research, National Cancer Institute) and E0771BR are a kind gift from Qing Chen (Wistar Institute) and were grown under the same conditions. For crystal violet staining, 5 x 10^4^ cells were plated and subjected to the treatments as described in the individual figures and then stained with 0.5% crystal violet prepared in a 1:1 methanol-water solution followed by PBS washes. Cells stably overexpressing OGT (GeneCopoeia, EX-Z3428-M13–10-OGT plasmid), CDK5 (GeneCopoeia, EX-A0348-Lv120), HA-ACSS2-WT, HA-ACSS2-S267D (GeneCopoeia, pReceiver), and HA-E2F-1-pRcCMV was a gift from William Kaelin (Addgene plasmid # 21667) were generated through production of lentivirus. All cell lines were quarterly tested for mycoplasma (abmGood PCR Mycoplasma Detection Kit). Following thawing, cells were only used for experiments for up to ten passages.

### Production of Lentivirus and Viral Transduction

HEK-293T cells were grown to ∼90% confluency and transfected. Prior to transfection, 20 μg of shRNA or overexpression plasmid DNA, 10 μg VSVG, 5 μg RSV, and 5 μg RRE were gently mixed and incubated in 1.5 mL of Optimem for 5 minutes. Concurrently, 105 μl of PEI was added dropwise to 1.5 mL of Optimem and incubated for 5 minutes. Following the 5 minute incubation, the PEI solution was added dropwise to the DNA solution and incubated for at least 30 minutes. The PEI-DNA solution was then added dropwise to the HEK-293T cells already plated with 5 mL of Optimem, and the cells were incubated overnight in the transfection media. Approximately 16-18 hours later, the transfection media was replaced with normal growth media. Viral supernatants were collected at 24 and 48 hours following the media change. These supernatants were passed through a 0.45 μm filter and portioned into 1 mL aliquots to be stored at −80°C if not for immediate use. 1 mL aliquots from each collection time point where mixed with 2mL of growth media and 1:500 8mg/mL polybrene and added to 75% confluent cell line of interest for 6 hours and replaced with 10mL of growth media followed by the appropriate antibiotic selection.

### RNA interference

Control shRNA was acquired from Addgene (plasmid 1864), from D. Sabatini (Massachusetts Institute of Technology). Control-scrambled shRNA sequence used was: CCTAAGGTTAAGTCGCCCTCGCTCTAGCGAGGGCGACTTAACCTT. All other shRNA constructs were acquired from Sigma and shRNA sequence used: for OGT-1, GCCCTAAGTTTGAGTCCAAATCTCGAGATTTGGACTCAAACTTAGGGC, and for OGT-2, GCTGAGCAGTATTCCGAGAAACTCGAGTTTCTCGGAATACTGCTCAGC. Cdk5 shRNA sequence used: for Cdk5-1, CCGGGTGAACGTCGTGCCCAAACTCCTCGAGGAGTTTGGGCACGACGTTCACTTTTTTG, and for Cdk5-2, CCGGCAGAACCTTCTGAAGTGTAACCTCGAGGTTACACTTCAGAAGGTTCTGTTTTTTG. ACSS2 shRNA sequence used: CCGGGCTTCTGTTCTGGGTCTGAATCTCGAGATTCAGACCCAGAACAGAAGCTTTTTG. Human E2F1 shRNA (Sigma): TAACTGCACTTTCGGCCCTTT. Mouse E2F1 shRNA (Sigma): CCATACCCTCTGTCCCAATAG.

### Animal models of cancer

Nu/Nu athymic 4-6 week old mice from Charles River Laboratories (Wilmington, MA, USA) were immobilized using the Just for Mice ^TM^ Stereotaxic Frame (Harvard Apparatus, Holliston, MA, USA) and injected intracranially with 5 μl of a 20,000 cells/μl solution of MDA-MB-231, MDA-MB-231BR cells stably expressing luciferase and pReceiver-WT-ACSS2 or pReceiver-S267D-ACSS2, N-Flag-OGT, HA-CDK5 overexpression or containing either Control or ACSS2, CDK5, OGT shRNA. For mammary fat pad (MFP) tumors, female athymic nude Nu/Nu mice (Charles River, Wilmington, MA, USA) (4–6 weeks old) were anesthetized with 4% isoflurane and inoculated with 1.5 x 10^6^ MDA-MB-231 cells stably expressing luciferase and either pReceiver-WT-ACSS2 or pReceiver-S267D-ACSS2 and containing either Control or ACSS2 shRNA. Mice were injected intraperitoneally with 200 µl of D-Luciferin solution (30 mg/ml; Caliper Life Sciences, Hopkinton, MA) and results analyzed using Living Image software (Caliper Life Sciences, Waltham, MA, USA). For intracranial injections, mice were euthanized 3 weeks after injection. Brains were dissected, fixed in 4% formalin and prepared for processing/sectioning/embedding/blocking to generate paraffin-embedded slides. For MFP injections, tumors were measured weekly along and perpendicular to the longest dimension using digital calipers (Fowler Co., Inc., Newton, MA, USA). Tumor volumes were calculated as V=(length)x(width)2 x0.52. After 8 weeks, tumors were excised, weighed, and photographed. All protocols involving the use of animals were approved by the Institutional Animal Care and Use Committee at Drexel University College of Medicine.

### Human samples

Paraffin embedded brain metastatic carcinoma tissue microarrays (GL861a) containing 43 cases of metastatic carcinoma from different primary tumors were purchased from Biomax. Slides were subjected to immunohistochemistry and stained for O-GlcNAc and pACSS2. Each slide was given a score from 1 (low) −3 (high) based on the observed intensity of each staining. Final pACSS2 expression was determined as a ratio of pACSS2 to its correspondent ACSS2 staining score.

### Reagents

Anti-actin (Santa Cruz Biotechnology; Dallas, TX, USA), anti-OGT, anti-ACSS2, anti-Cdk5, (Cell Signaling Technology; Danvers, MA, USA), anti-O-GlcNAc (Sigma Aldrich; St. Louis, MO, USA) Anti-E2F1, Anti-GPX4, Anti-SLC7A11 (Abcam). Puromycin, Polybrene, crystal violet (Sigma Aldrich, St. Louis, MO, USA). Rabbit polyclonal antibody recognizing phosphorylated ACSS2 S267 was created by as previous described [16]. D-luciferin potassium salt (Perkin Elmer, Waltham, MA, USA). Plasmids used were pReceiver N-Flag OGT, pReceiver HA-ACSS2, HA-ACSS2-S267A, and HA-ACSS2-S267D (Genecopoeia; Rockville, MD, USA). Cdk5-HA was a gift from Sander van den Heuvel (Addgene plasmid # 1872). D-luciferin potassium salt (Perkin Elmer, Waltham, MA, USA), and crystal violet (Sigma Aldrich, St. Louis, MO, USA).

### mRNA expression

RNA was extracted using Trizol (Invitrogen) as per manufacturer’s instruction from MDA-MB-231BR cells Taqman gene expression assay primer probes for ACSS2 (Hs01122829_m1), GAPDH (Hs02758991_g1) E2F1 (Hs00153451_m1) SLC7A11 (Hs00921938_m1) and GPX4 (Hs00989766_g1) were purchased from Thermofisher Scientific. qRT-PCR was performed using Brilliant II qRT-PCR Master Mix 2 Kit (Agilent, San Diego, CA, USA) and analysis was performed using Applied Biosystems 7500 machine. Briefly, isolated RNA concentrations were determined by measuring them on a NanoDrop, and all samples were diluted to match the sample with the lowest concentration. A mastermix was then generated for the individual primers using 8.8 μl dH O, 12.5 μl Brilliant PCR master mix, 0.4 μl diluted dye (0.5 μl dye into 250 μl dH O), 0.1 μl reverse transcriptase, and 1.25 μl primer per sample. A total reaction volume of 25 μl was used by gently mixing 2 μl of sample RNA and 23 μl of the mastermix into the reaction tubes, carefully avoiding bubbles. The samples were then loaded into the Applied Biosystems 7500 machine and the experimental setup was as follows: acquire experiment as “Quantitation ΔΔCT”, targets and samples were assigned, the initial holding stage adjusted to 50°C for 30 minutes, followed by a secondary holding stage at 95°C for 10 minutes, then 40 cycles consisting of 95°C for 15 seconds and cooling to 60°C for 1 minute were completed. Data were exported from the Applied Biosystems software and expression levels were analyzed using Data Assist v2.0 (Life Technologies, Grand Island, NY, USA).

### Immunoblotting

Immunoblotting protocols have been previously described. Briefly, cell lysates from 1-5 x 10^6^ cells were prepared in radioimmune precipitation assay buffer (150 mM NaCl, 1% NP40, 0.5% DOC, 50 mM Tris HCL at pH 8, 0.1% SDS, 10% glycerol, 5 mM EDTA, 20 mM NaF, and 1 mM Na_3_VO_4_) supplemented with 1 μg/ml each of pepstatin, leupeptin, aprotinin, and 200 μg/ml PMSF. Lysates were cleared by centrifugation at 16,000 x *g* for 15 minutes at 4 °C and analyzed by SDS-PAGE and autoradiography with chemiluminescence. Proteins were analyzed by immunoblotting using primary antibodies indicated above.

### RNA-Seq

RNA was extracted using Trizol (Invitrogen) as per manufacturer’s instruction from MDA-MB-231BR cells stably expressing shSCR or shACSS2 (3 biological replicates per condition). RNA-Seq and analysis was performed by BGI Tech.

### Immunohistochemical staining

The slides containing brain metastatic tumors were deparaffinized by Xylene and subsequent rehydration was done by decreasing concentrations of ethanol-water mixture. Antigen retrieval was done by citrate buffer immersion and steaming the slides for 45 minutes. Tissue was treated with 200-400 ul 1% BSA+5% serum PBS solution for 1hr. Primary incubation was done at 4° C overnight using OGT, O-GlcNAc RL2 (Santa Cruz 1:50), p-S267 ACSS2 (1:100), ACSS2 (Cell Signaling Technology 1:100), GFAP (Life Technologies 2.2B10 1:100) antibodies. Secondary antibody incubation was done for an hour at RT with respective antibodies at 1: 100 dilution. For IHC the stain was developed using the DAB kit by Vectastain (Vector Labs, Burlingame, CA, USA). Finally, slides were mounted and imaged under a light microscope. The staining patterns of tumor tissues of GBM were assessed by a board-certified pathologist. For immunofluorescence, alexa fluor conjugated secondary antibodies were used at 1:100 dilution in the dark, slides were mounted and imaged under a fluorescent microscope. H&E and Ki-67 staining were performed at TJU, Department of Pathology, Philadelphia, PA.

### Clonogenic survival Assay

Cells were cultured and detached in their respective growth media to assess clonogenic survival, then counted using a hemocytometer. 1,000 cells were plated in triplicate in 6-well plates in their respective growth media and allowed to form colonies. Cells were grown for 14 days. Upon conclusion of the growth period, colonies were stained with crystal violet. Colony numbers were quantified based on number of colonies with >40 cells.

### BODIPY Staining of Cells

Cells were stained with 5μM BOPIDY C11 (Invitrogen) in PBS for 25 min at 37C in the dark and washed in 1xPBS prior to imaging on EVOS FL (Life Technologies) using GFP.

### Flow Cytometry

Cells were prepared according to manufacturer protocol (BD Pharmingen, Propidium Iodine) for cell death and BODIPY C11 (Invitrogen) for lipid peroxidation. Briefly, cells were trypsinized (0.25% Trypsin), counted, washed twice with 1xPBS, and resuspended in 100uL 1X binding buffer incubated with 5 uL Propidium Iodine (PI) staining or 5uM BODIPY solution for 15 minutes in the dark at room temperature. Following incubation, the volume was brought up to 500uL of 1X binding buffer. Tubes were then analyzed using a Guava easyCyte flow cytometer. All data were collected and analyzed using a Guava EasyCyte Plus system and CytoSoft (version 5.3) software (Millipore). Data are gated and expressed relative to the appropriate unstained and single-stained controls.

### *Ex vivo* brain slice model

Organotypic hippocampal cultures were prepared as described previously. Briefly, adult mice (6-8 week) or mice after 12 days following intracranial injection were decapitated and their brains rapidly removed into ice-cold (4°C) sucrose-aCSF composed of the following (in mM): 280 sucrose, 5 KCl, 2 MgCl2, 1 CaCl2, 20 glucose, 10 HEPES, 5 Na^+^-ascorbate, 3 thiourea, 2 Na^+^-pyruvate; pH=7.3. Brains were blocked with a sharp scalpel and sliced into 250 µm sections using a McIlwain-type tissue chopper (Vibrotome inc). Four to six slices were placed onto each 0.4 µm Millicell tissue culture insert (Millipore) in six-well plates, 1 ml of medium containing the following: Neurobasal medium A (Gibco), 2% Gem21-Neuroplex supplement (Gemini), 1% N2 supplement (Gibco), 1% glutamine (Invitrogen), 0.5% glucose, 10 U/ml penicillin, and 100 μg/ml streptomycin (Invitrogen), placed underneath each insert. One-third to one-half of the media was changed every 2 days. Tumor growth was monitored via bioluminescence imaging on the IVIS 200 system (Perkin Elmer) and results analyzed using Living Image software (Caliper Life Sciences, Waltham, MA, USA). For MTS assay, individual brain slices were transferred to a 96-well plate and subjected to Promega CellTiter 96® Aqueous One Solution (Cat: G3582) mixed in a 1:5 ratio with culture media and treated as previously described. Tissues were incubated at 37(, 5% CO_2_ for 4 hours and absorbance at 490nm was measured with Tecan Spark Microplate reader.

### Statistical Analysis

All results shown are results of at least three independent experiments and shown as averages and presented as mean ± s.e. P-values were calculated using a Student’s two-tailed test (* represents p-value ≤ 0.05 or **p-value ≤ 0.01 or as marked in figure legend). Statistical analysis of growth rate of mice was performed using ANCOVA. *p-value < 0.05.

## Acknowledgements

We thank Anna Ramesh and Aman Gupta for their technical assistance. This work was supported by NIH-NCI grant UO1CA244303 (to MJR), PA Breast Cancer Coalition Award (to MJR), and Coulter-Drexel Translational Research Award (to AD & MJR).

## Author contributions

M.J.R., L.C., E.M.E., R.G.Y. conceived and planned all experiments. E.M.E., R.G.Y., L.C., N.A., J.M., M.K. performed all *in vitro* experiments. E.M.E., L.C., R.G.Y., J.M., A.N.T. performed all *in vivo* experiments. L.C., W.G. performed immunohistochemistry. A.D. provided AD-5584. M.J.R., E.M.E., R.G.Y., L.C. interpreted results. M.J.R., L.C. E.M.E. wrote first draft of manuscript. M.J.R., E.M.E., L.C., R.G.Y., A.D., N.L.S. edited manuscript.

## Supplemental Figures

**Supplemental Figure 1. (A)** Representative images of immunohistochemical staining for H&E, O-GlcNAc and p-S267-ACSS2 on coronal sections from mice inoculated with MDA-MB-231 or MDA-MB-231BR cells. **(B)** Immunohistochemical staining for O-GlcNAc, ACSS2 and pACSS2 on a brain metastatic carcinoma tissue microarray (n=86) (4X scale bar 1000 μm) and representative images of pACSS2 scoring as low, medium or high based on a 1, 2 and 3 score respectively for pACSS2 and O-GlcNAc staining. **(C)** Cell lysates from MDA-MB-231 cells stably overexpressing control or OGT RNA were collected for immunoblot analysis with the indicated antibodies. **(D)** Representative images of bioluminescent (top) detection of tumors from mice injected with Control and OGT MDA-MB-231 cells 14 days post-injection. Representative images of H&E analysis (bottom) on coronal sections from mice harboring Control or OGT overexpression RNA tumors at Day 14. Bioluminescence signal from mice injected with MDA-MB-231 cells over expressing Control (n=3) or OGT mice (n=3) (bottom). Student’s t-test reported as mean ± SEM; *p<0.05. **(E)** Cell lysates from MDA-MB-231 cells stably overexpressing control or CDK5 RNA were collected for immunoblot analysis with the indicated antibodies. **(F)** Representative images of bioluminescent (top) detection of tumors from mice injected with Control and CDK5 MDA-MB-231 cells 14 days post-injection. Representative images of H&E analysis (bottom) on coronal sections from mice harboring Control or CDK5 overexpression RNA tumors at Day 14. Bioluminescence signal from mice injected with MDA-MB-231 cells over expressing Control (n=3) or OGT mice (n=3) (bottom). Student’s t-test reported as mean ± SEM; *p<0.05.

**Supplemental Figure 2. (A)** Cell lysates from 4T1BR cells stably expressing shRNA against Scramble or ACSS2 were collected for immunoblot analysis with the indicated antibodies. **(B)** Representative images of bioluminescent detection of tumors from mice injected with shSCR and shACSS2 4T1BR cells 14 days post-injection. Bioluminescence signal from mice injected with 4T1BR expressing shRNA against Scramble (n=3) or ACSS2 mice (n=3) (top). Student’s t-test reported as mean ± SEM; *p<0.05. Representative images of brain sections stained for H&E 14-days-post treatment (bottom). **(C)** Measurement of acetyl-CoA extracted from MDA-MB-231 cells stably expressing Control, Wildtype ACSS2 and ACSS2 S267D mutant. Student’s t-test reported as mean ± SEM. * = p-value < 0.05.

**Supplemental Figure 3. (A)** Cell lysate from MDA-MB-231BR stably expressing shRNA against Scramble, or CDK5 and overexpressing HA-ACSS2-S267D7 were collected for immunoblot analysis with the indicated antibodies. **(B)** Clonogenic survival assay of MDA-MB- 231BR cells stably expressing shRNA against control, CDK5 with orHA-ACSS2-267D overexpression. Student’s t-test reported as mean ± SEM. * = p-value < 0.05. **(C)** Representative images of bioluminescent (top) detection of tumors from mice injected with shSCR, shCDK5 and S267D-shCDK5 MDA-MB-231BR luciferase expressing cells 21 days post-injection. Representative images of H&E on coronal sections from mice harboring tumors at Day 21 (bottom).

**Supplemental Figure 4. (A)** *Ex vivo* brain slices (without tumor) were treated for 6 days with ACSS2 inhibitor (VY-3-249) with either DMSO or ACSS2 inhibitor collected on day 6, and analyzed for cell viability (MTS assay). As a positive control, slices were treated with paraformaldehyde (PFA) for 2 hrs, rendering the brain slices non-viable (n=3). Student’s t-test reported as mean ± SEM; *p<0.05. **(B)** Quantification of propidium iodine staining flow cytometry of 4T1BR cells treated with DMSO, ACSS2 inhibitor (VY-3-249), or ACSS2i + Ferrostatin-1 (Fer-1). Data reported as One-way ANOVA with Tukey’s multiple comparisons test reported as mean ± SEM *p-value < 0.05, **p<0.01. **(C)** Quantification of lipid peroxides with Bopidy C11 and flow cytometry of 4T1BR cells treated with DMSO, ACSS2 inhibitor (VY-3-249), or ACSS2i + Ferrostatin-1 (Fer-1). Data reported as One-way ANOVA with Tukey’s multiple comparisons test reported as mean ± SEM **p-value < 0.01, ***p-value < 0.001. **(D)** Cell lysates from 4T1BR cells stably expressing shRNA against Scramble or ACSS2 were collected for immunoblot analysis with the indicated antibodies. **(E)** Quantification of propidium iodine staining flow cytometry of 4T1BR cells expressing shRNA against Scramble or ACSS2, treated with and without Fer-1. Data reported as One-way ANOVA with Tukey’s multiple comparisons test reported as mean ± SEM *p-value < 0.05. **(F)** Quantification of lipid peroxides with Bopidy C11 and flow cytometry of 4T1BR cells stably expressing shRNA against Sramble or ACSS2 treated with and without Fer-1. Data reported as One-way ANOVA with Tukey’s multiple comparisons test reported as mean ± SEM **p-value < 0.01. **(G)** Representative images depicting tumor growth in organotypic brain slices derived from mice intracranially injected with MDA-MB-231BR-luc cells detected via bioluminescence. Brain slices containing tumors are treated with Control (DMSO), Erastin, and Ferrostatin-1 (Fer-1) for indicated days.

**Supplemental Figure 5. (A)** MDA-MB-231BR cells expressing shSCR or shACSS2 were subjected to RNAseq. Volcano plot showing up and downregulated genes. **(B)** Enrichment plot demonstrating E2F1 genes negatively enriched in shACSS2. **(C)** Quantification of propidium iodine staining flow cytometry of MDA-MB-231BR cells stably overexpressing control or HA- E2F1 treated with ACSS2 inhibitor (VY-3-249) for 48 hours. Data reported as One-way ANOVA with Tukey’s multiple comparisons test reported as mean ± SEM *p-value < 0.05, **p<0.01. **(D)** Cell lysates from parental MDA-MB-231, 4T1 **(E)**, E0771 **(F)** and brain trophic cells were collected for immunoblot analysis with the indicated antibodies.

**Supplemental Figure 6. (A)** Cell lysates collected from MDA-MB-231BR cells stably overexpressing control or HA-E2F1 were collected for immunoblot analysis with the indicated antibodies. **(B)** Total RNA was collected from MDA-MB-231BR cells stably expressing shRNA against Scramble or ACSS2. Quantification of qRT-PCR performed on RNA extracts analyzing E2F1, SLC7A11, and GPX4 gene expression normalized to GAPDH. Two-way ANOVA reported as mean ± SEM.* p-value < 0.05. **(C)** Cell lysates of MDA-MB-231BR cells treated with ACSS2 inhibitor (VY-3-249) 50 μM for 48 hours were collected for immunoblot analysis with the indicated antibodies. **(D)** Cell lysates of 4T1BR cells treated with ACSS2 inhibitor (VY-3-249) 50 μM for 48 hours were collected for immunoblot analysis with the indicated antibodies.

**Supplemental Figure 7. (A)** Cell lysates from 4T1BR cells treated with control (DMSO) or ACSS2 inhibitor AD-5584 (100 μM) for 48 hrs were collected for immunoblot analysis with the indicated antibodies. (**B)** Quantification of propidium iodine staining flow cytometry of 4T1BR cells treated with DMSO, AD-5584, or AD-5584 + Fer1. Data reported as One-way ANOVA with Tukey’s multiple comparisons test reported as mean ± SEM *p-value < 0.05, **p<0.01. **(C)** Quantification of lipid peroxides with Bopidy C11 and flow cytometry of 4T1BR cells treated with DMSO, AD-5584, or AD-5584 + Fer1. Data reported as One-way ANOVA with Tukey’s multiple comparisons test reported as mean ± SEM **p-value < 0.01. Representative images depicting tumor growth in organotypic brain slices derived from mice intracranially injected with 4T1BR-lcu cells detected via bioluminescence. Brain slices containing tumors are treated with Control (DMSO), ACSS2 inhibitor (AD-5584) or AD-5584 and Ferrostatin-1 (Fer-1) for indicated days (top). **(E)** Representative images of bioluminescent detection of tumors from Nu/Nu mice injected with luciferase tagged MDA-MB-231BR cells at Day 0 (prior to drug treatment) and at 14 days post-drug treatment. Data are quantified and presented as average Relative Bioluminescence signal from mice injected with MDA-MB-231BR cells treated with Vehicle (n=3) or AD-5584 treated mice (n=3) (right). Student’s t-test reported as mean ± SEM; *p<0.05. **(F)** Quantification of weights (grams) of mice injected with 4T1BR cells and treated with vehicle (n=3) or AD-5584 (n=3) for 10 days, analyzed with two-way ANOVA, n.s. **(G)** Quantification of weights (grams) of mice injected with MDA-MB-231BR cells and treated with vehicle (n=3) or AD-5584 (n=3) for 14 days, analyzed with two-way ANOVA, n.s.

