## Supplemental Figures for "ACSS2 regulates ferroptosis in an E2F1-dependent manner in breast cancer brain metastatic cells"

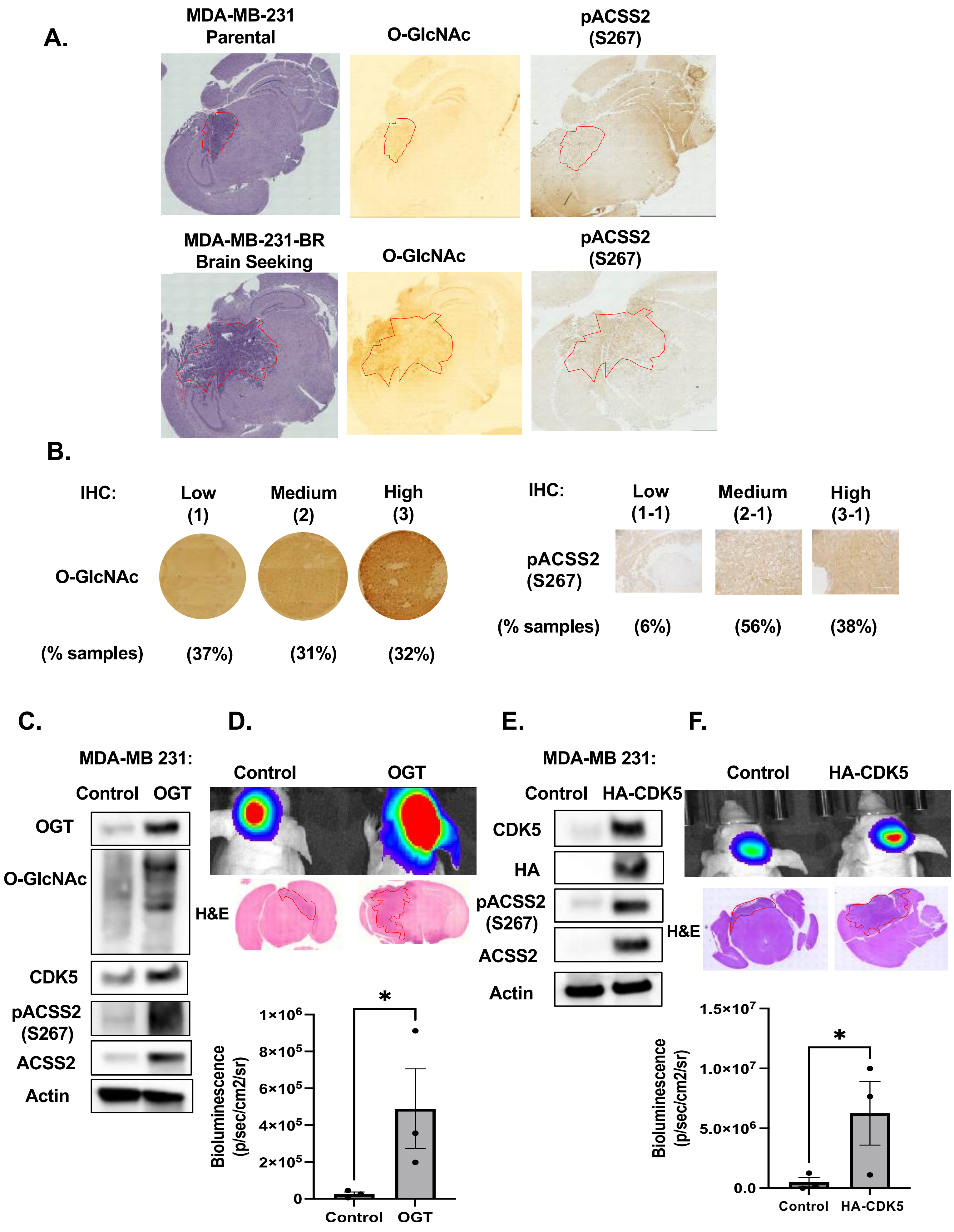
A.
MDA-MB-231
Parental
O-GlcNAc
pACSS2
(S267)
MDA-MB-231-BR
Brain Seeking
O-GlcNAc
pACSS2
(S267)
B.
IHC:
Low
(1)
Medium
(2)
High
(3)
O-GlcNAc
(% samples)
(37%)
(31%)
(32%)
IHC:
Low
(1-1)
Medium
(2-1)
High
(3-1)
pACSS2
(S267)
(% samples)
(6%)
(56%)
(38%)
C.
MDA-MB 231:
Control
OGT
OGT
O-GlcNAc
CDK5
pACSS2
(S267)
ACSS2
Actin
D.
Control
OGT
H&E
Bioluminescence
(p/sec/cm2/sr)
1×10⁶
8×10⁵
6×10⁵
4×10⁵
2×10⁵
0
*
Control
OGT
E.
MDA-MB 231:
Control
HA-CDK5
CDK5
HA
pACSS2
(S267)
ACSS2
Actin
F.
Control
HA-CDK5
H&E
Bioluminescence
(p/sec/cm2/sr)
1.5×10⁷
1.0×10⁷
5.0×10⁶
0.0
*
Control
HA-CDK5

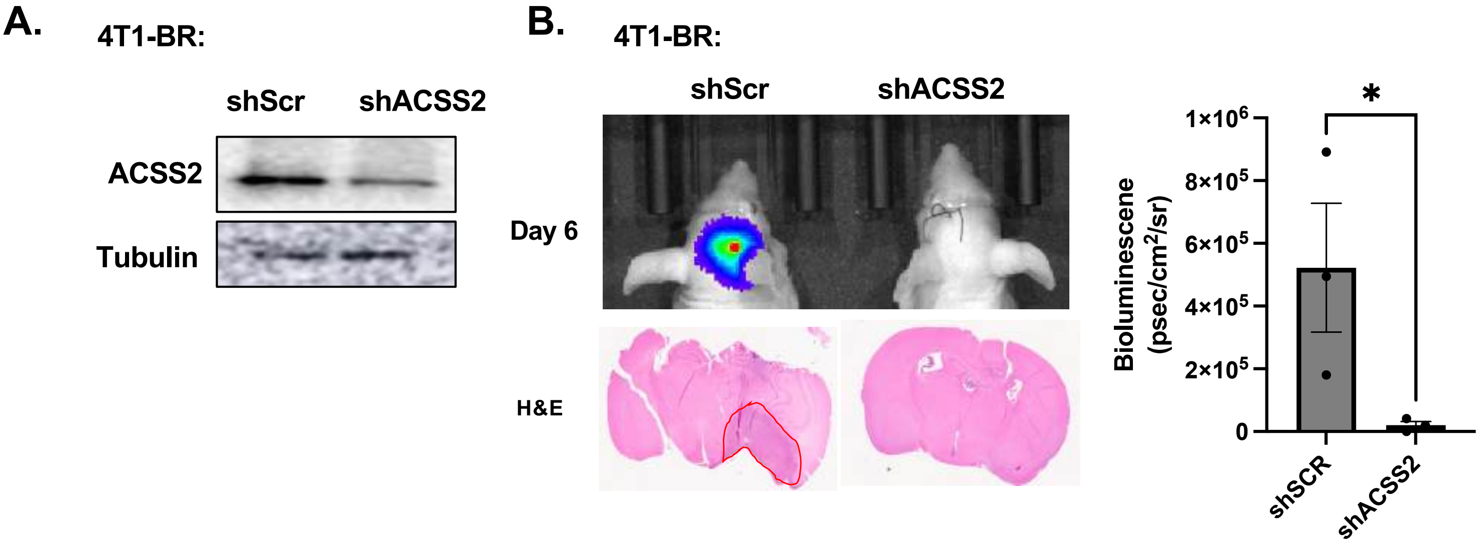
A.
4T1-BR:
shScr
shACSS2
ACSS2
Tubulin
B.
4T1-BR:
shScr
shACSS2
Day 6
H&E
*
1×10^6
8×10^5
6×10^5
4×10^5
2×10^5
0
Bioluminescene
(psec/cm^2/sr)
shSCR
shACSS2

Esquea et al. Supplemental Figure 3

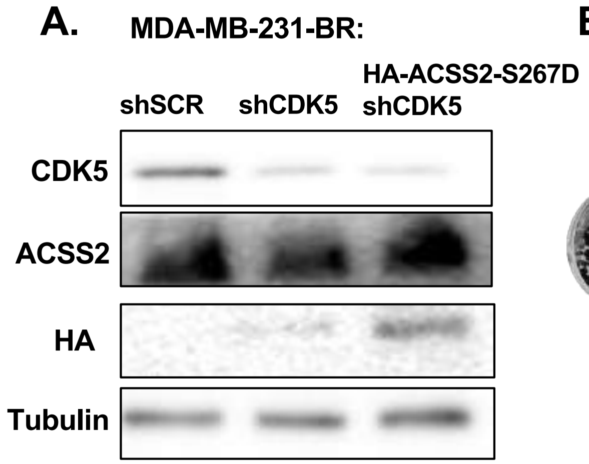


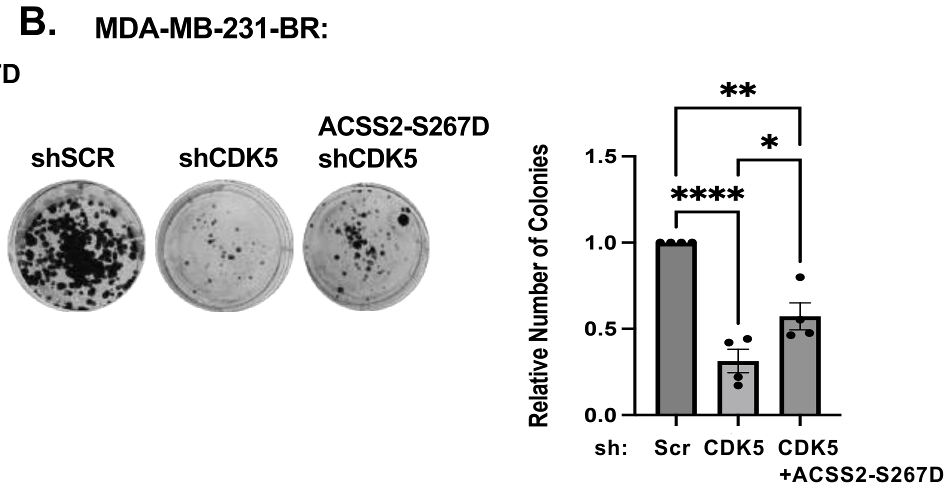


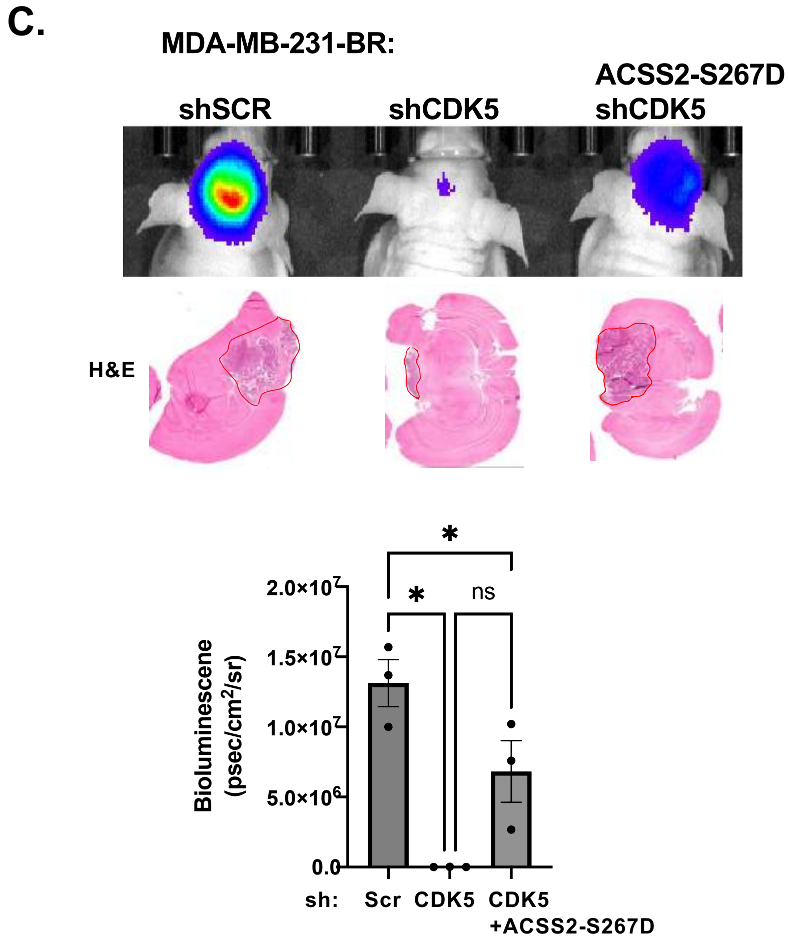

Esquea et al. Supplemental Figure 4

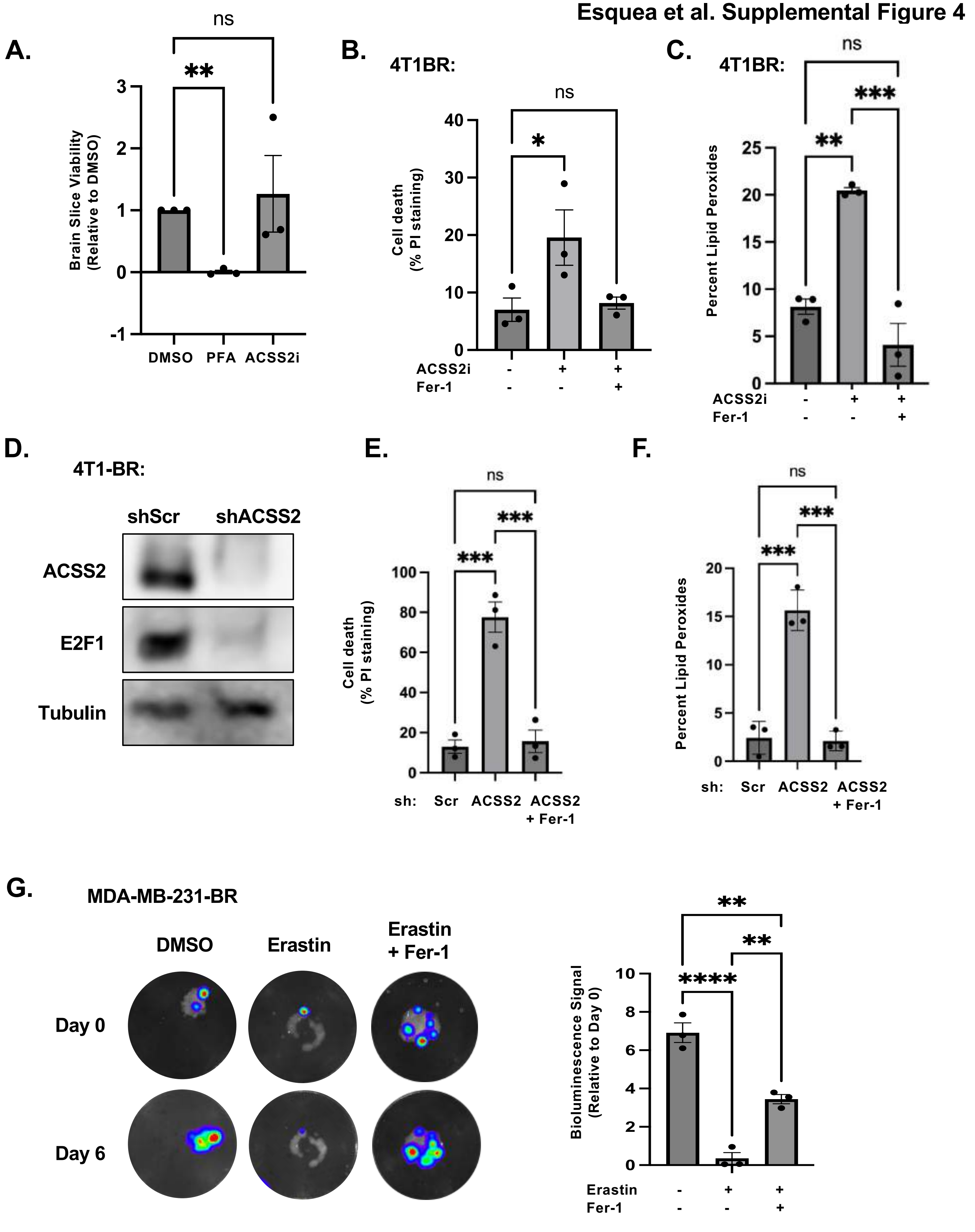

Esquea et al. Supplemental Figure 5

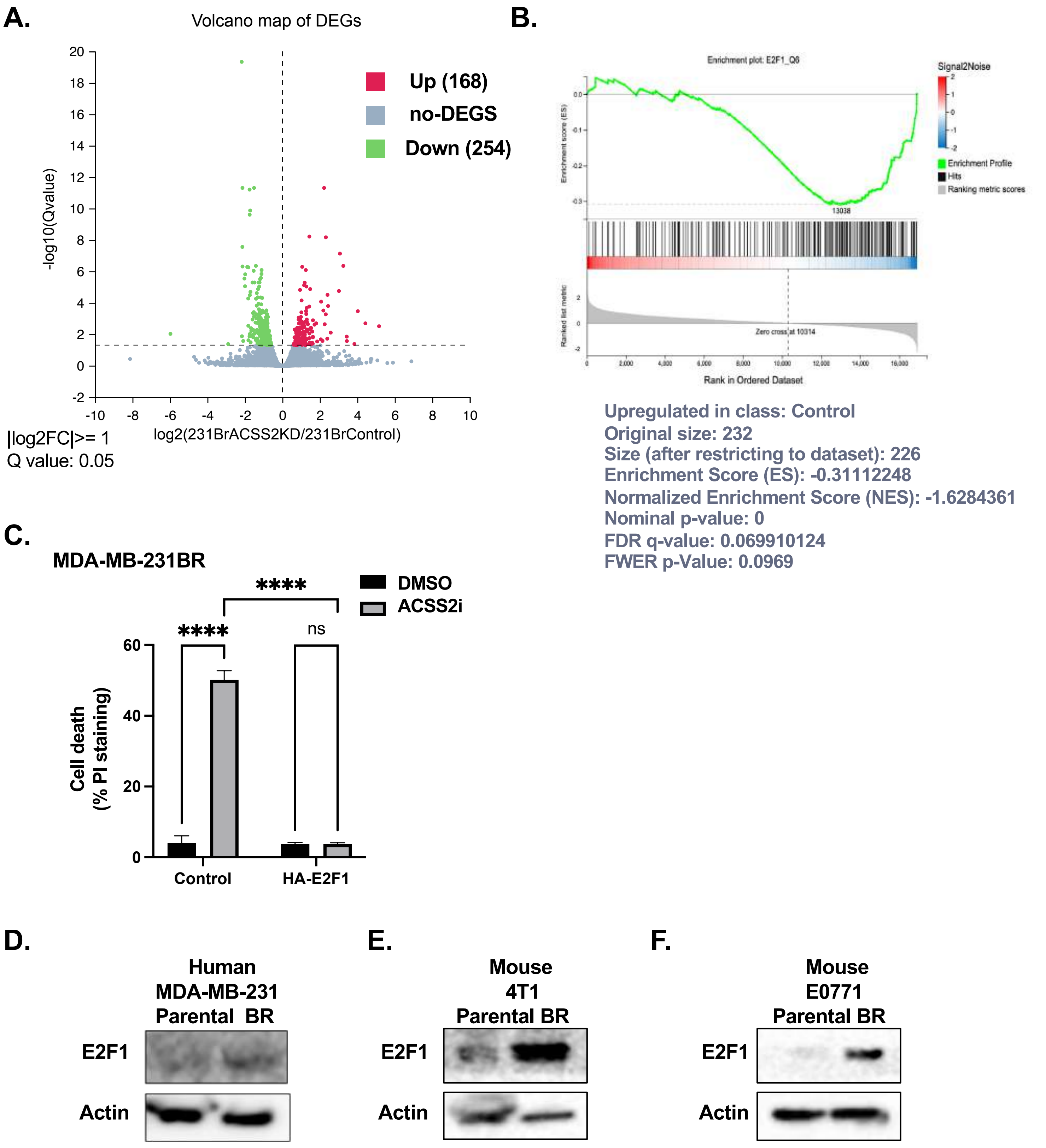

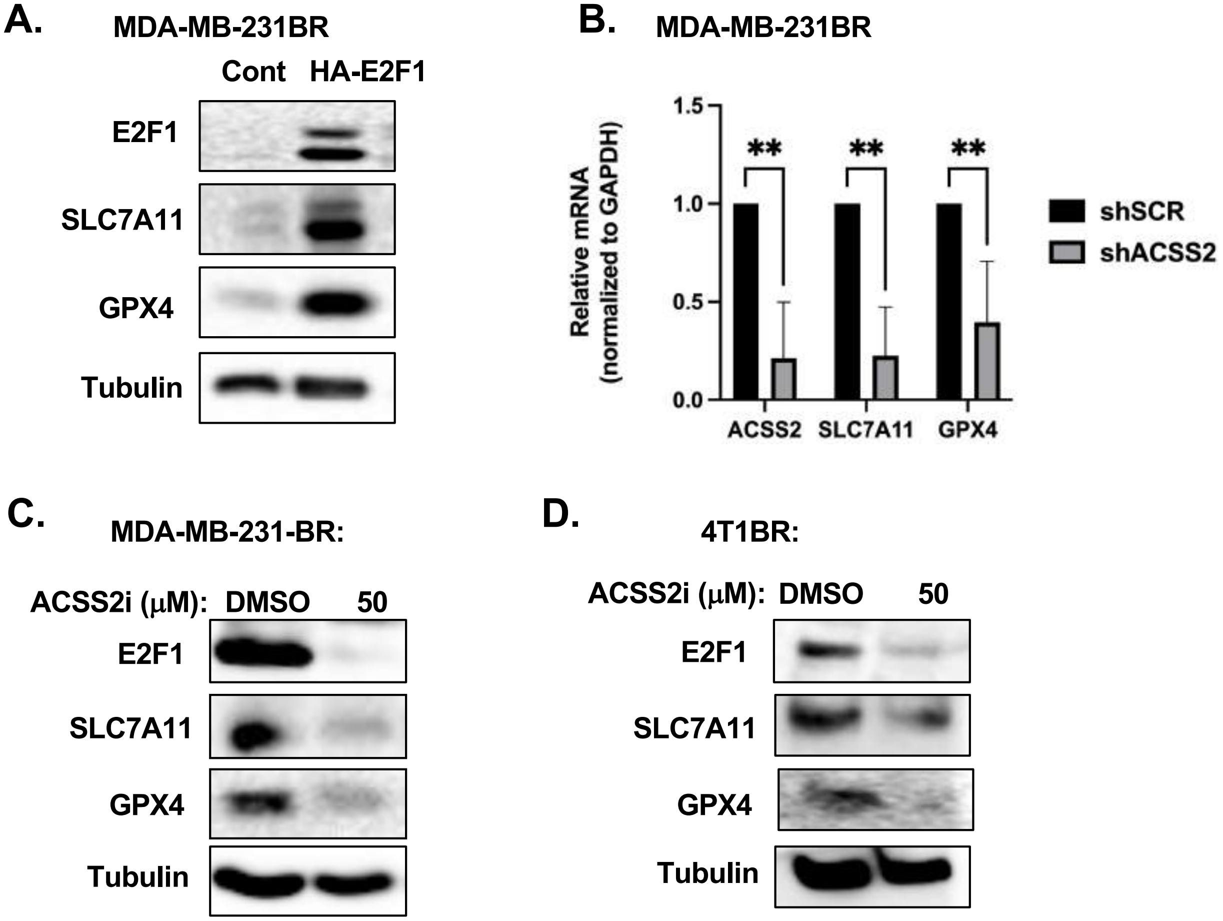
A.
MDA-MB-231BR
Cont
HA-E2F1
E2F1
SLC7A11
GPX4
Tubulin
B.
MDA-MB-231BR
Relative mRNA
(normalized to GAPDH)
1.5
1.0
0.5
0.0
**
**
**
ACSS2
SLC7A11
GPX4
shSCR
shACSS2
C.
MDA-MB-231-BR:
ACSS2i (μM):
DMSO
50
E2F1
SLC7A11
GPX4
Tubulin
D.
4T1BR:
ACSS2i (μM):
DMSO
50
E2F1
SLC7A11
GPX4
Tubulin

Esquea et al. Supplemental Figure 7

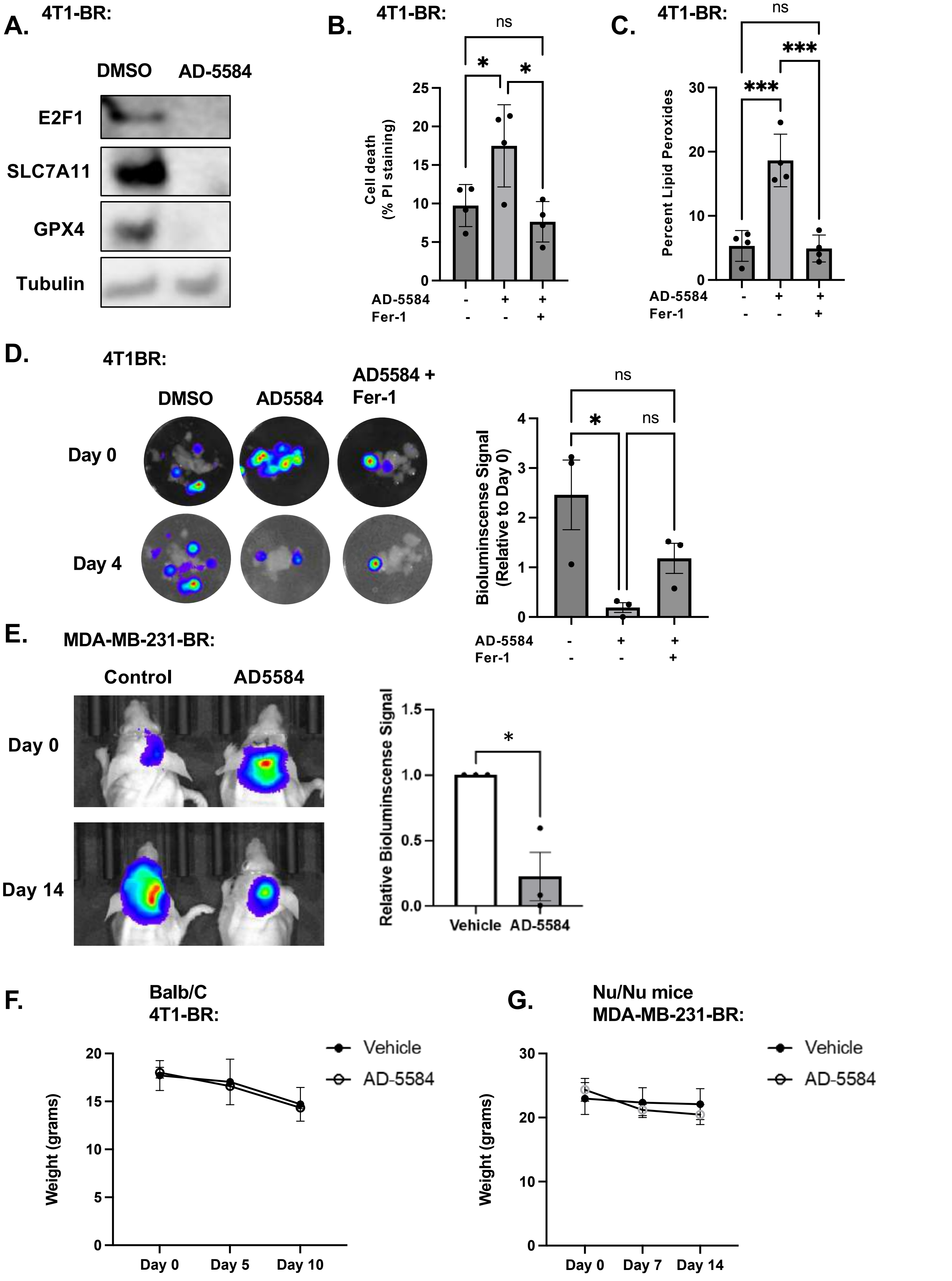
